# Optimized but not maximized cue integration for 3D visual perception

**DOI:** 10.1101/611087

**Authors:** Ting-Yu Chang, Byounghoon Kim, Lowell Thompson, Adhira Sunkara, Raymond Doudlah, Ari Rosenberg

## Abstract

Reconstructing three-dimensional (3D) scenes from two-dimensional (2D) retinal images is an ill-posed problem. Despite this, our 3D perception of the world based on 2D retinal images is seemingly accurate and precise. The integration of distinct visual cues is essential for robust 3D perception in humans, but it is unclear if this mechanism is conserved in non-human primates, and how the underlying neural architecture constrains 3D perception. Here we assess 3D perception in macaque monkeys using a surface orientation discrimination task. We find that perception is generally accurate, but precision depends on the spatial pose of the surface and available cues. The results indicate that robust perception is achieved by dynamically reweighting the integration of stereoscopic and perspective cues according to their pose-dependent reliabilities. They further suggest that 3D perception is influenced by a prior for the 3D orientation statistics of natural scenes. We compare the data to simulations based on the responses of 3D orientation selective neurons. The results are explained by a model in which two independent neuronal populations representing stereoscopic and perspective cues (with perspective signals from the two eyes combined using nonlinear canonical computations) are optimally integrated through linear summation. Perception of combined-cue stimuli is optimal given this architecture. However, an alternative architecture in which stereoscopic cues and perspective cues detected by each eye are represented by three independent populations yields two times greater precision than observed. This implies that, due to canonical computations, cue integration for 3D perception is optimized but not maximized.

**Author summary:** Our eyes only sense two-dimensional projections of the world (like a movie on a screen), yet we perceive the world in three dimensions. To create reliable 3D percepts, the human visual system integrates distinct visual signals according to their reliabilities, which depend on conditions such as how far away an object is located and how it is oriented. Here we find that non-human primates similarly integrate different 3D visual signals, and that their perception is influenced by the 3D orientation statistics of natural scenes. Cue integration is thus a conserved mechanism for creating robust 3D percepts by the primate brain. Using simulations of neural population activity, based on neuronal recordings from the same animals, we show that some computations which occur widely in the brain facilitate 3D perception, while others hinder perception. This work addresses key questions about how neural systems solve the difficult problem of generating 3D percepts, identifies a plausible neural architecture for implementing robust 3D vision, and reveals how neural computation can simultaneously optimize and curb perception.

## Introduction

Three-dimensional (3D) visual perception is a significant achievement of the primate brain [1]. Because the eyes detect two-dimensional (2D) projections of the world, like a movie on a screen, 3D structure must be estimated. Creating 3D percepts from 2D images is a nonlinear optimization problem plagued by ambiguities and noise [2]. Human perceptual [3–5] and neuroimaging [6–9] studies show that integrating distinct visual cues resolves ambiguities and improves 3D estimates. The neural implementation of optimal cue integration is, theoretically, a linear process [10], but nonlinear computations such as quadratic nonlinearities and divisive normalization are also widely implicated in neural processing [11–17]. Because such nonlinearities reduce the independence of neuronal stimulus representations, they can conceivably impose limits on the precision of perception. We tested this hypothesis using psychophysics and computational modeling to evaluate how non-human primates (NHPs) perceptually integrate two visual cues which have prominent roles in human 3D vision: stereoscopic and perspective cues.

Stereoscopic cues arise from comparisons of left and right retinal images, which differ because the eyes are horizontally offset [18, 19]. Perspective cues originate from the projection of the 3D world onto 2D retinae [20, 21]. The reliability of the 3D information carried by these cues depends on an object’s spatial pose (i.e., position and orientation) [4, 5]. Specifically, stereoscopic cue reliability decreases with distance (Fig 1A) and perspective cue reliability increases with slant (Fig 1B). Human studies reveal that the integration of these cues is weighted according to their reliabilities [4, 5], but little is known about how NHPs perceptually integrate these cues.

**Fig 1.**
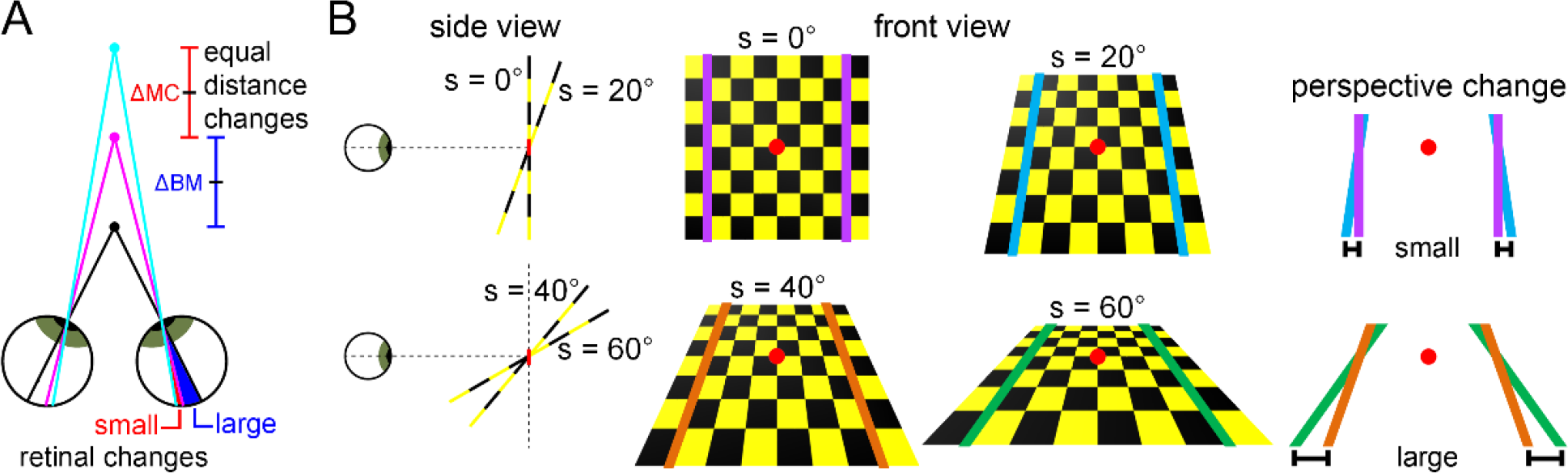
3D cue reliabilities depend on object pose. **(A)** Stereoscopic cue reliability decreases with distance. Equivalent changes in distance result in smaller retinal image changes at greater distances. The distance between the black and magenta dots (ΔBM) is equal to the distance between the magenta and cyan dots (ΔMC), but the retinal change is larger for ΔBM than ΔMC. **(B)** The reliability of perspective cues increases with orientation in depth (slant). The rate at which parallel lines converge in a 2D projection increases with slant. This is illustrated with a checkerboard rotated about the horizontal axis passing through the red dot. Colored lines are parallel in the world. A 20° slant (s) rotation produces a smaller perspective change between slants of 0° and 20° (top row) than slants of 40° and 60° (bottom row).

Using an eight-alternative forced choice (8AFC) tilt discrimination task, we quantified how perception depends on planar surface pose. Contributions of stereoscopic and perspective cues to perception were evaluated using cue-isolating and combined-cue stimuli. For stereoscopic cues, performance decreased with distance from the fixation plane, consistent with geometric limitations of stereovision and the physiology of stereopsis [22]. For both cues, performance increased with slant. We further found evidence of a 3D analogue of the ‘oblique effect’ (more accurate and precise perception of cardinal than oblique tilts) [23–27], consistent with the influence of a prior for the 3D orientation statistics of natural scenes [28, 29].

Perception of combined-cue stimuli was consistent with an optimal integration strategy [30], with the cues dynamically reweighted according to their pose-dependent reliabilities. We found that perception was well explained by a neural architecture in which stereoscopic and perspective cues are represented by independent populations, with perspective signals from the two eyes combined via a quadratic nonlinearity and divisive normalization prior to their integration with stereoscopic cues. Cue integration was optimal given this architecture (population responses were linearly summed [10]). However, an alternative architecture in which stereoscopic as well as left and right eye perspective cues are all represented independently yielded ~2 times greater precision. This indicates that 3D perception is optimized, but not maximized, and suggests that the precision of 3D perception is curbed by nonlinear canonical computations in the representation of perspective cues. Analogous limitations may exist for other sensory processes with multiple inputs signaling the same cue types, as occurs in audition, vestibular processing, and bimanual touch. Our findings suggest that cue integration is a conserved mechanism by which primates achieve robust 3D vision, and that the co-occurrence of multiple canonical computations (linear summation, quadratics, and divisive normalization) simultaneously optimizes and curbs perception.

## Results

### Accuracy and precision of combined-cue 3D perception

The 3D orientation of a planar surface can be described by two angular variables: tilt and slant [31, 32]. Tilt specifies the direction that the plane is oriented in depth (e.g., top-near), and slant specifies how much it is oriented in depth (i.e., its steepness; Fig 1B). We trained two rhesus macaques to perform an 8AFC tilt discrimination task. The monkeys reported a plane’s tilt with an eye movement to a choice target (Fig 2A). Slant and distance were varied to evaluate how tilt perception changed with the plane’s pose. We first quantified perception of combined-cue stimuli defined by stereoscopic and perspective cues (Fig 2B and 2C).

**Fig 2.**
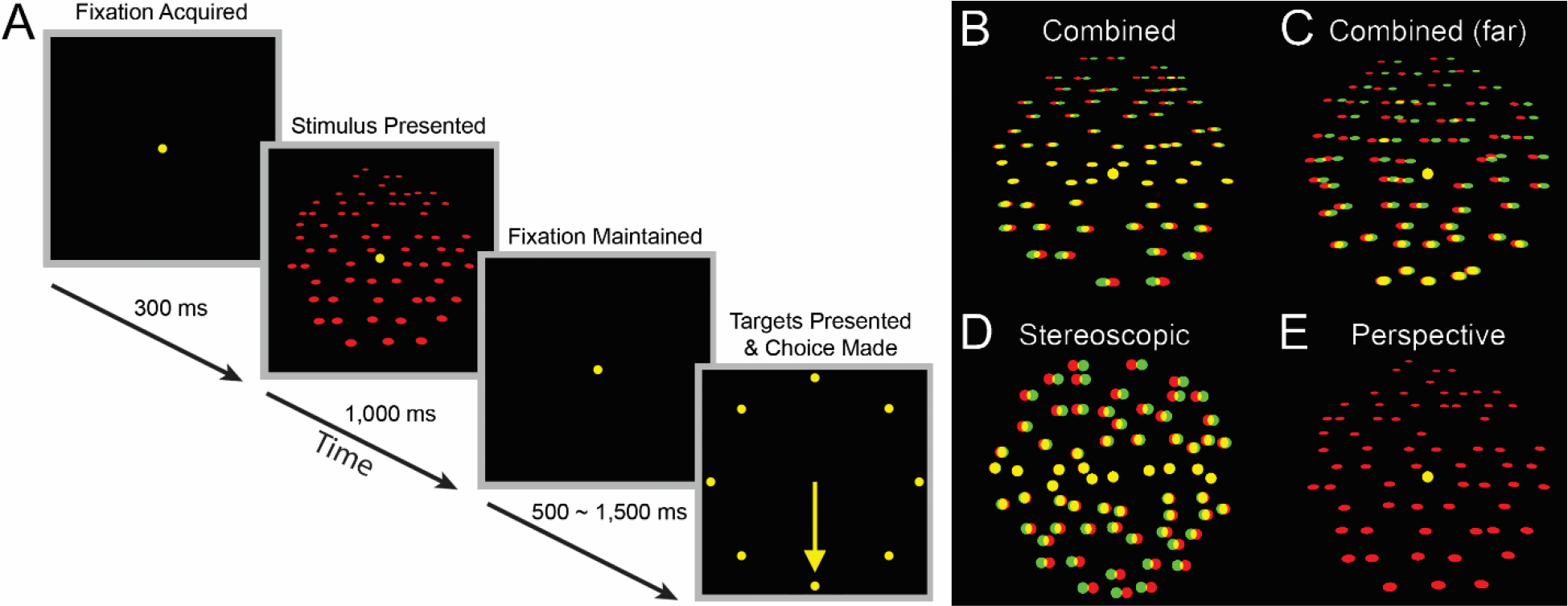
Discrimination task and stimuli. **(A)** Eight alternative tilt discrimination task. Fixation was maintained on a target for 300 ms. A plane centered on the target was then presented for 1,000 ms. Fixation was then held for an additional 500-1,500 ms before the target disappeared and eight choice targets appeared. The tilt of the plane was reported through a saccade to the corresponding choice target. For example, the bottom target for a bottom-near (tilt = 270°) plane. **(B-E)** Example planes (tilt = 270°, slant = 60°). For clarity, dot size is exaggerated and dot number reduced from the actual experiments. Stimuli are illustrated as red–green anaglyphs. **(B)** Combined-cue stimulus at 57 cm (fixation distance). **(C)** Combined-cue stimulus at 77 cm (all dots behind fixation). **(D)** Stereoscopic cue stimulus at 57 cm. **(E)** Perspective cue stimulus at 57 cm (left eye presentation).

Representative data characterizing tilt perception for combined-cue stimuli are shown in Fig 3. These data show error distributions of reported tilts (ΔTilt = Reported Tilt − Presented Tilt) calculated using all 8 tilts for 12 combinations of slant and distance. Perception was quantified in terms of accuracy (i.e., if there were systematic deviations between perceived and actual tilts) and precision (i.e., the variability of the percepts). Accuracy and precision were quantified using the bias (μ) and concentration (*κ*) parameters of von Mises probability density function fits (equation 1, see **Methods**) to the error distributions [33]. No bias (μ = 0°) indicates perfect accuracy, and larger values of *κ* (taller and narrower densities) indicate greater precision.

**Fig 3.**
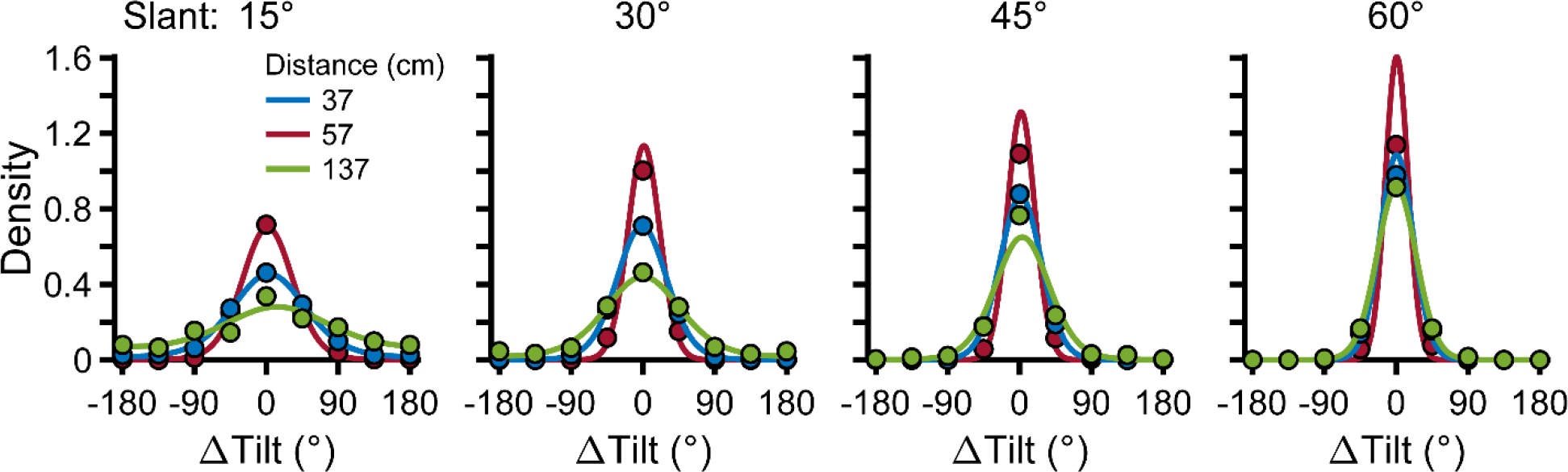
Tilt discrimination with combined-cue stimuli. Each curve shows a probability density function describing the errors in reported tilts made by Monkey L with combined-cue stimuli, calculated using all 8 tilts. Each column shows densities at a given slant, and colors correspond to different distances. Correct choices correspond to ΔTilt = 0°. The probability that an error of a given ΔTilt was made is shown with a point. Solid curves are von Mises density fits used to quantify the accuracy and precision of perception. All curves peaked close to ΔTilt = 0°, indicating that performance was accurate. Taller and narrower densities indicate greater precision. Precision increased monotonically with slant (curves grow taller from left to right), and showed an inverted U-shape as a function of distance (red curves are taller than blue and green curves).

#### Accuracy results

All of the density functions shown in Fig 3 peaked close to 0°, indicating that tilt perception was accurate. Indeed, across the 24 slant–distance combinations tested with Monkey L, the biases were centered close to 0° and narrowly distributed: circular mean μ = 1.50°, circular standard deviation (SD) = 3°. The results were similar for Monkey F: mean μ = 0.54°, SD = 2.33° (N = 32 slant–distances). We repeated this analysis at each tilt individually, and again found little bias (S1A Fig). These results indicate that tilt perception with combined-cue stimuli was accurate over a wide range of poses defined by distance, slant, and tilt.

Although the biases at each tilt were small, Monkey L showed an overall pattern consistent with an oblique effect for planar tilt. Across all cardinal tilts, the median absolute bias was 3.66° (N = 24 slant–distances × 4 tilts = 96), but 8.21° across all oblique tilts. Consistent with the influence of a prior for cardinal tilts, which are more frequent than oblique tilts in natural scenes [28, 29], the oblique biases were significantly larger than the cardinal biases (circular median test, *p* = 5.32×10^−4^). However, for Monkey F, the median absolute biases at cardinal (3.31°, N = 32 × 4 = 128) and oblique (3.55°) tilts were not significantly different (*p* = 0.80). Individual differences in the strength of the 2D oblique effect are similarly observed in humans [24, 25].

#### Precision results

The precision of combined-cue tilt perception depended on surface pose in two ways. First, precision increased monotonically with slant, as seen in the right marginals of Fig 4. This is also evident in Fig 3 by comparing the density functions across columns. With increasing slant (left to right in the figure), the densities became taller and narrower (larger *κ*). Second, precision showed an inverted U-shape as a function of distance, as shown in the top marginals of Fig 4. Likewise, this is seen in Fig 3, where the density functions at 57 cm (maroon curves) are taller and narrower than those at 37 cm (blue) or 137 cm (green). How precision depended on both slant and distance is summarized using heat maps in Fig 4. These precision landscapes reflect the interaction of the monotonic relationship between precision and slant and the inverted U-shape relationship between precision and distance, resulting in more gradual decreases in precision with distance at larger slants. Precision peaked at the largest slant (60°) and ~20 cm behind the plane of fixation for both monkeys. Although precision varied with surface pose, performance was above chance at all slant–distance combinations (Rayleigh test for circular uniformity, all *p* ≤ 4.96×10^−14^ and significant after correcting for 24 or 32 comparisons for Monkeys L and F, respectively).

**Fig 4.**
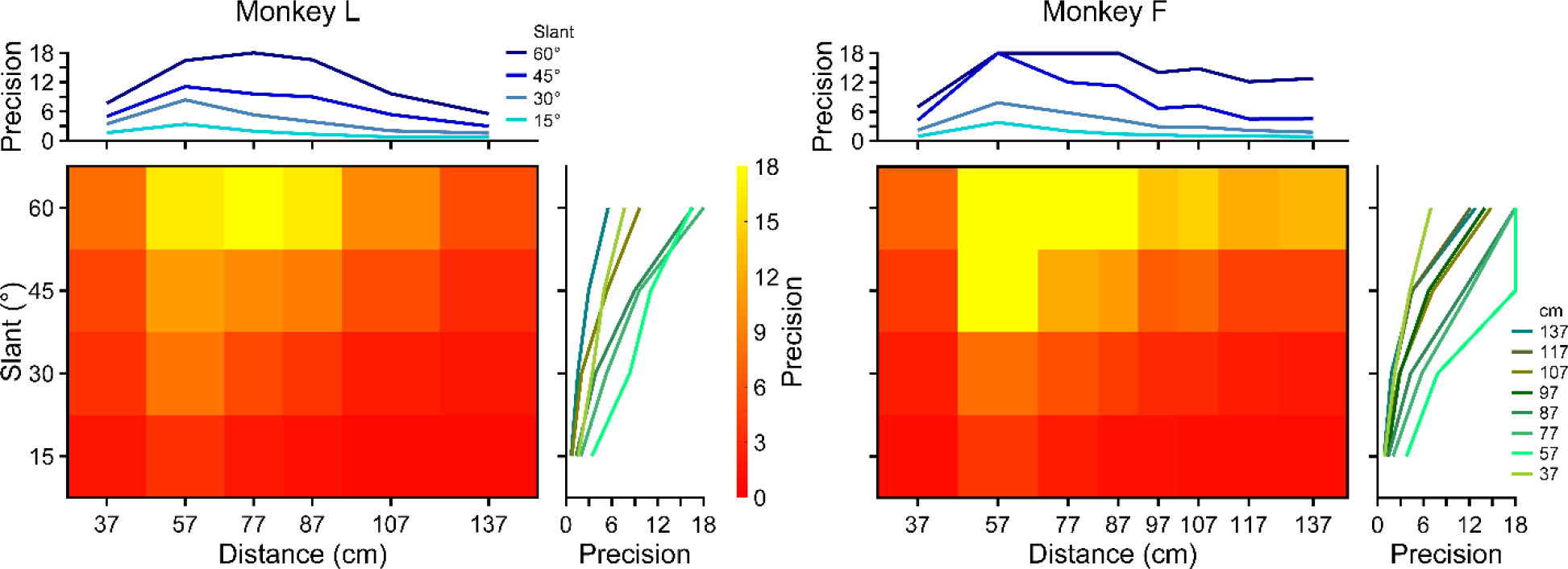
Precision of tilt perception with combined-cue stimuli. Heat maps showing the precision (*κ*) of tilt perception as a function of slant and distance for Monkeys L (left) and F (right), calculated using all 8 tilts. Red hues indicate lower precision and yellow hues indicate higher precision. Precision peaked at the largest tested slant (60°) and 20 cm behind the plane of fixation (57 cm). The larger the slant, the slower performance fell off with distance, giving a wedge-shaped appearance to the precision landscapes. Right marginal curves show *κ* as a function of slant for each distance. Precision increased monotonically with slant. Upper marginal curves show *κ* as a function of distance for each slant. Precision showed an inverted U-shape as a function of distance. An upper bound of 18 was set on *κ* (see **Methods**).

We further found that precision did not differ significantly as a function of tilt for either monkey (S2A Fig), and that the results generalized to larger stimuli (S3A Fig). However, similar to the bias results, we found that Monkey L showed an oblique effect when we grouped precisions at cardinal and oblique tilts. For Monkey L, the median precision at cardinals tilts (6.95; N = 96) was significantly larger than at oblique tilts (4.78), Mann-Whitney U test (*p* = 5.99×10^−3^). For Monkey F, the median precisions at cardinal (4.99, N = 128) and oblique (5.01) tilts were not significantly different (*p* = 0.64). These results parallel findings from human perceptual studies which indicate that the precision of 3D perception depends on the pose-dependent reliabilities of the available visual cues [4, 5], and further suggest that there are individual differences in the extent to which the 3D orientation statistics of natural scenes impact perception.

### Contributions of stereoscopic cues to 3D perception

Next, we assessed tilt perception using stimuli that isolated stereoscopic cues (Fig 2D). Control experiments confirmed that the stimuli contained no perspective cues that could be used to perform the task, and that performance was unaffected by a potential stereoscopic–perspective cue conflict [5] (S4 Fig). Error distributions of reported tilts were again calculated using all 8 tilts, and the accuracy and precision of perception were quantified using von Mises fits.

#### Accuracy results

For both monkeys, mean stereoscopic cue biases across all slant–distance combinations were again close to 0°, indicating that perception was generally accurate (Monkey L: mean μ = −3.04°, SD = 19.64°, N = 24; Monkey F: mean μ = 1.87°, SD = 20°, N = 32). However, the biases were broadly distributed. Examination of the biases at individual tilts suggested that this variability was due to geometric factors (S1B Fig). At surface poses with low stereoscopic cue reliability (i.e., combinations of large distances and small slants) precision was particularly poor, and the biases were correspondingly large. In contrast, performance was accurate at poses where the precision was reasonably high. Thus, perception was accurate so long as the cues were sufficiently reliable for the monkeys to perform the task well.

For the stereoscopic cue stimuli, Monkey L once again showed a pattern of biases consistent with an oblique effect. Across all cardinal tilts, the median absolute bias was 7.76° (N = 96), but 15.58° across all oblique tilts (circular median test, *p* = 3.89×10^−3^). Monkey F did not show this pattern: the median absolute biases at cardinal (8.19°, N = 128) and oblique (8.26°) tilts were not significantly different (*p* = 1). However, when precision was low, both monkeys showed a bias towards reporting bottom-near (270°) tilts (S1B Fig). This bias is consistent with the influence of a prior for ground planes, which are preponderant in natural scenes [28, 29].

#### Precision results

Precision landscapes over slant and distance are shown for the stereoscopic cue stimuli in Fig 5A. The overall patterns resembled the combined-cue landscapes (Monkey L: r = 0.96, *p* = 3.37×10^−13^; Monkey F: r = 0.81, *p* = 1.49×10^−8^). There was a monotonic relationship between precision and slant, indicating that stereoscopic cue reliability increases with slant. There was also an inverted U-shape relationship between precision and distance which is explained by geometric and physiological factors. The falloff in precision with distance is consistent with the decreasing reliability of stereoscopic cues (Fig 1A). The falloff in precision with distance from the fixation plane (both toward and away from the monkey) is consistent with the limited range of horizontal disparities represented by the visual system [22]. While the stereoscopic and combined-cue precision landscapes were similar in pattern, precision was significantly lower for the stereoscopic cue stimuli than the combined-cue stimuli (Wilcoxon signed-rank test; Monkey L: *p* = 1.82×10^−5^, N = 24; Monkey F: *p* = 7.95×10^−7^, N = 32). Indeed, at combinations of large distances and small slants, performance with stereoscopic cue stimuli was not significantly different from chance (outlined in black in Fig 5A; Rayleigh test for circular uniformity, corrected for multiple comparisons). At greater distances, performance was at chance levels even with larger stimuli (S3B Fig). Thus, stereoscopic cues did not contribute to tilt perception beyond ~137 cm (less for small slants), indicating that perspective cues mediated above chance performance with combined-cue stimuli at those poses (Fig 4 and S3A Fig).

**Fig 5.**
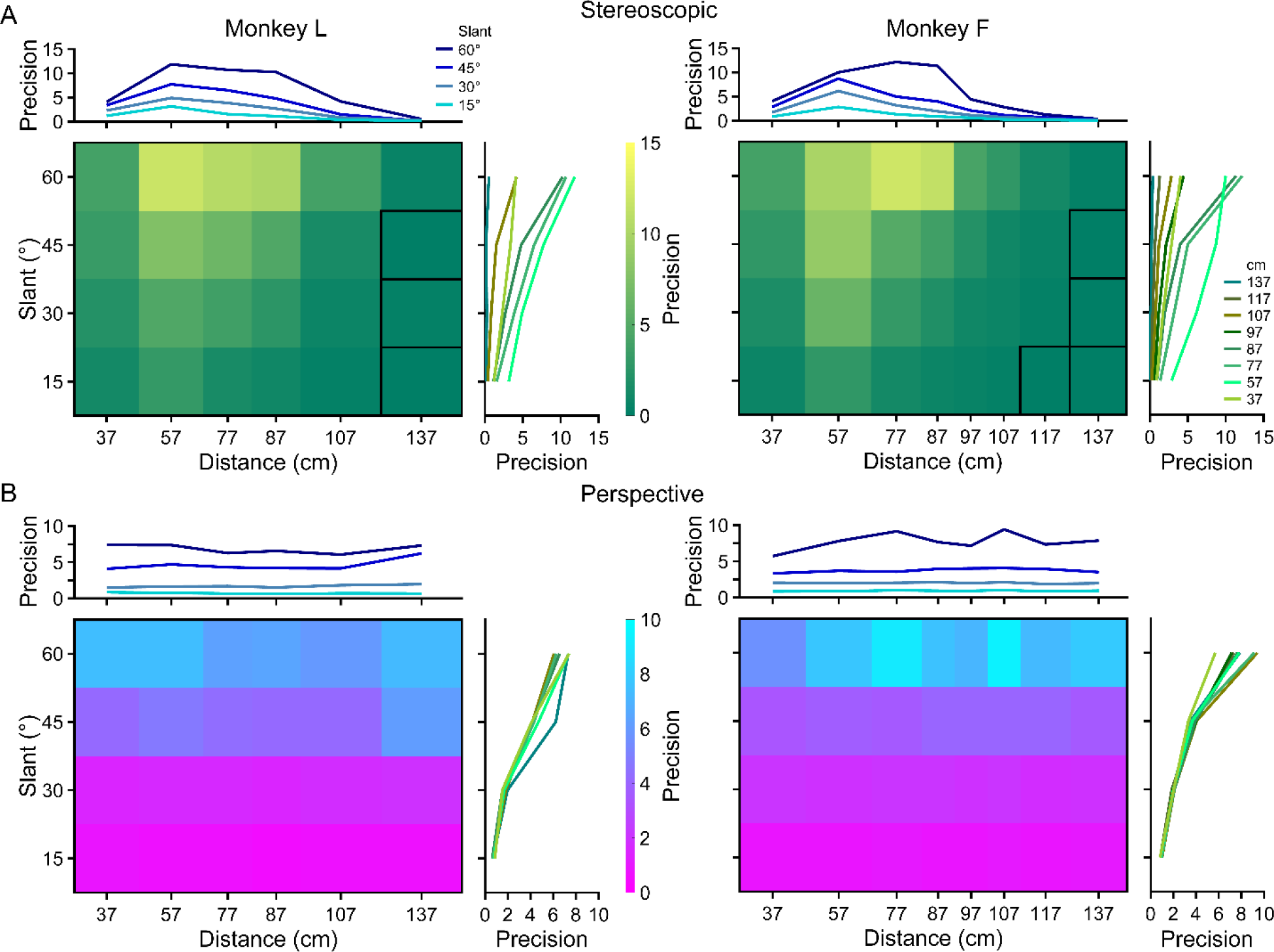
Precision of tilt perception with cue-isolating stimuli. **(A)** Stereoscopic cue stimuli. Precision (*κ*) increased monotonically with slant, and showed an inverted U-shape as a function of distance. Performance was at chance levels for combinations of small slants and large distances (outlined in black). **(B)** Perspective cue stimuli. Precision increased monotonically with slant, and was largely independent of distance. Plotted as in Fig 4.

We further found that precision did not differ significantly as a function of tilt (S2B Fig), or between cardinal (median *κ* = 3.46 and 1.80 for Monkeys L and F, respectively) and oblique (median *κ* = 3.18 and 1.76 for Monkeys L and F, respectively) tilts for either monkey (Mann-Whitney U test; Monkey L: *p* = 0.37, N = 96; Monkey F: *p* = 0.99, N = 128). Together, these results indicate that the contributions of stereoscopic cues to 3D perception are constrained by a combination of viewing geometry and physiology.

### Contributions of perspective cues to 3D perception

Next, we assessed tilt perception using stimuli that isolated perspective cues (Fig 2E). To eliminate stereoscopic cues, we presented single eye views of combined-cue stimuli to the appropriate eye, and only the fixation target to the other eye. Performance was comparable with the two eyes (S5 Fig), so responses to left and right eye stimulus presentations were pooled together. Error distributions of reported tilts were calculated using all 8 tilts, and the accuracy and precision of tilt perception were quantified using von Mises fits.

#### Accuracy results

The perspective cue biases were centered close to 0° and narrowly distributed across all slant–distance combinations (Monkey L: mean μ = 0.31°, SD = 3.04°, N = 24; Monkey F: mean μ = 2.19°, SD = 2.69°, N = 32), indicating that perception was accurate irrespective of the surface pose. Indeed, there was little bias at any individual tilt (S1C Fig). Although the biases at each tilt were small, Monkey L showed an overall pattern of biases consistent with an oblique effect for planar tilt: the median absolute bias at oblique tilts (8.71°, N = 96) was significantly larger than the median absolute bias at cardinal tilts (4.10°), circular median test (*p* = 1.50×10^−3^). Monkey F did not show this pattern: median absolute biases at oblique (6.27°, N = 128) and cardinal (5.41°) tilts were not significantly different (*p* = 0.32). These results indicate that perspective cues support accurate perception of 3D orientation across a wide range of surface poses, and further suggest that there are individual differences in the extent to which the 3D orientation statistics of natural scenes influence perception based on perspective cues.

#### Precision results

Precision landscapes over slant and distance are shown for the perspective cue stimuli in Fig 5B. At all poses, performance was above chance. There was a monotonic relationship between precision and slant (greater precision at higher slants), consistent with the slant-dependent reliability of perspective cues (Fig 1B). Precision was independent of distance, reflecting that the perspective cues in our stimuli signaled orientation but not distance due to the elimination of absolute size cues (see **Methods**). For both monkeys, precision was significantly lower for the perspective cue stimuli than the combined-cue stimuli (Wilcoxon signed-rank test; Monkey L: *p* = 7.48×10^−4^; Monkey F: *p* = 1.86×10^−6^). Across all poses, the perspective and stereoscopic cue precisions were not significantly different (Wilcoxon signed-rank test; Monkey L: *p* = 0.49; Monkey F: *p* = 0.15). However, the relative precisions for the two cue types were distance dependent. For distances at or just behind the fixation plane (57, 77, and 87 cm), precision was significantly higher with stereoscopic cues (Wilcoxon signed-rank test; Monkey L: *p* = 4.88×10^−4^, N = 12; Monkey F: *p* = 4.88×10^−3^, N = 12). For nearer (37 cm) and further distances (> 87 cm), precision was significantly lower with stereoscopic cues (Wilcoxon signed-rank test; Monkey L: *p* = 6.84×10^−3^, N = 12; Monkey F: *p* = 1.03×10^−4^, N = 20).

We further found that precision did not differ substantially as a function of tilt (S2C Fig), and that the results generalized to larger stimuli (S3C Fig). However, similar to the bias results, we found that Monkey L showed an oblique effect when we grouped precisions at cardinal and oblique tilts. For Monkey L, the median precision at cardinals tilts (4.32; N = 96) was significantly larger than at oblique tilts (3.12), Mann-Whitney U test (*p* = 0.011). For Monkey F, the median precisions at cardinal (2.77, N = 128) and oblique (3.0) tilts were not significantly different (*p* = 0.12). Together, these results indicate that both stereoscopic and perspective cues contribute to 3D perception within peripersonal space, and that perspective cues extend 3D perception beyond the range supported by stereoscopic cues.

### Perceptual cue integration

The previous sections showed that perception was more precise for combined-cue than cue-isolated stimuli, and that the relative precisions for cue-isolated stimuli were pose-dependent. Given these results, we next tested if the cues were integrated optimally. That is, if stereoscopic and perspective cues were dynamically reweighted according to their pose-dependent reliabilities to maximize the precision of combined-cue stimulus perception. To test this hypothesis, we used cue integration theory to derive optimal predictions of the combined-cue bias 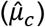 and precision 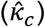 from the cue-isolated data (see **Methods**) [30]. We then compared the observed and optimal combined-cue biases and precisions to determine if the two cues were optimally reweighted on a trial-by-trial basis.

Representative error distributions and von Mises fits are shown for cue-isolated and combined-cue stimuli along with optimal predictions in Fig 6A–D. Observed (blue curves) and optimal (dashed black curves) combined-cue performances were highly similar. Across all slant– distance combinations, the observed and optimal biases were not significantly different from each other (circular median test for multiple samples, *p* = 0.13, N = 56, both monkeys). Likewise, the observed and optimal precisions were highly correlated (r = 0.94, *p* = 1.39×10^−27^, N = 56), distributed along the identity line (Fig 6E), and not significantly different from each other (Wilcoxon signed-rank test, *p* = 0.12, N = 56). The results were similar with larger stimuli (S3D Fig), and when cue integration was assessed separately for cardinal and oblique tilts (S6 Fig). These results suggest that, like humans, monkeys achieve robust 3D visual perception through the optimal integration of stereoscopic and perspective cues. Since all of the stimulus conditions were interleaved and presented pseudo-randomly, cue reweighting had to occur dynamically to match the vagaries of cue reliabilities that occurred with trial-to-trial changes in surface pose.

**Fig 6.**
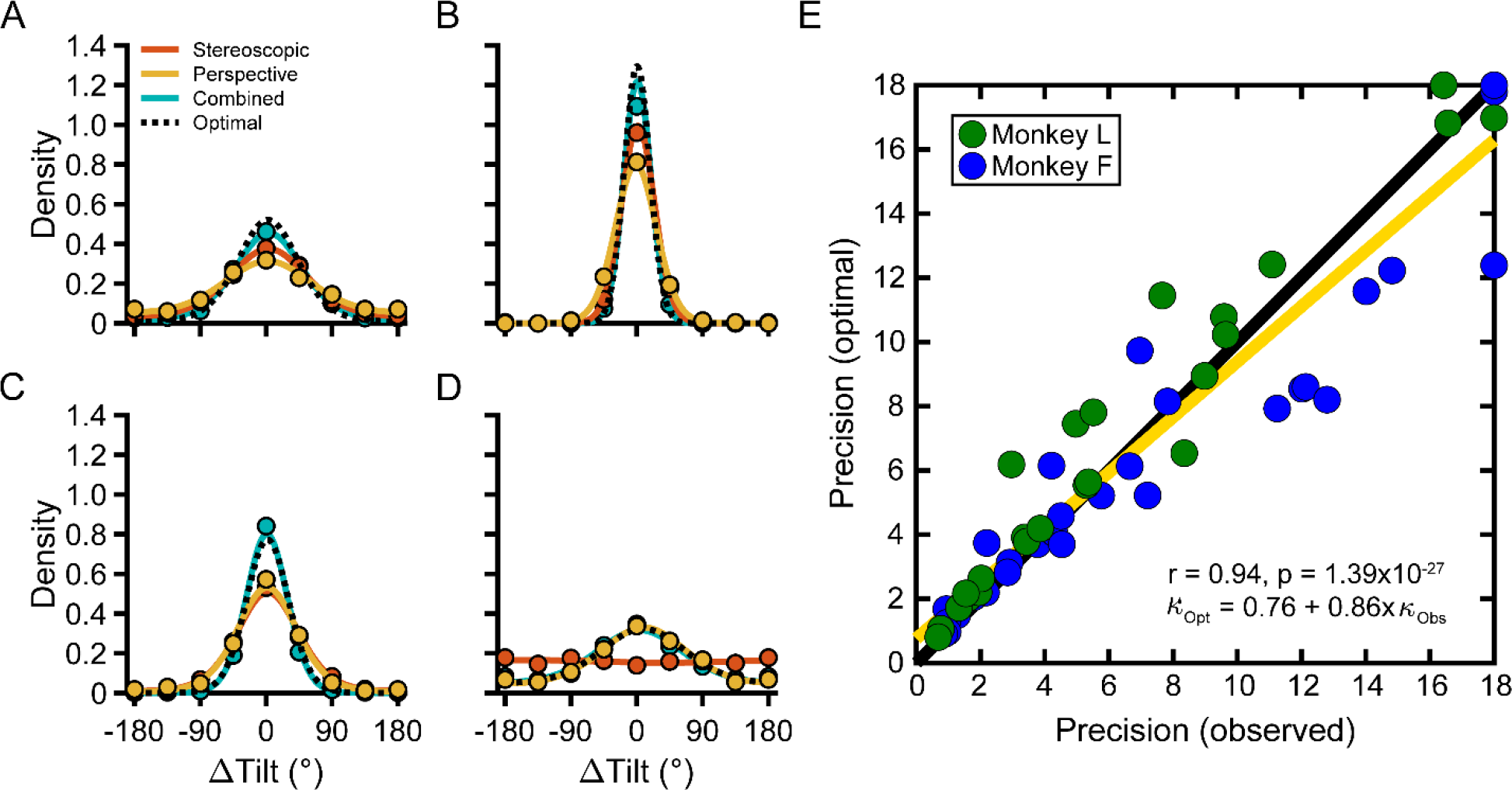
Optimal cue integration. **(A-D)** Representative densities. Solid curves show von Mises fits for each cue type (colors) and dotted back lines show optimal combined-cue performance. **(A)** Slant = 15°, distance = 77 cm (Monkey F). Stereoscopic *κ* > perspective *κ*. **(B)** Slant = 60°, distance = 87 cm (Monkey L). Stereoscopic *κ* > perspective *κ*. **(C)** Slant = 60°, distance = 107 cm (Monkey L). Stereoscopic *κ* ≈ perspective *κ*. **(D)** Slant = 45°, distance = 117 cm (Monkey F). Stereoscopic *κ* ≪ perspective *κ*. Combined-cue perception depended entirely on perspective cues. **(E)** Each point shows the optimal vs. observed combined-cue precision (*κ*) for a single slant–distance combination and monkey. The type-II regression line is shown in yellow (*κ* = 18 were excluded from the fit). Combined-cue precision was well predicted by optimal cue integration across a broad range of poses with different relative isolated-cue precisions.

### Neuronal models of 3D visual cue integration

We found optimal integration of stereoscopic and perspective cues, consistent with previous human results [4, 5]. However, previous studies did not consider that combined-cue stimuli actually contain three cues: stereoscopic, left eye perspective, and right eye perspective [34]. A distinction between left and right eye perspective cues may seem surprising, but the two retinal projections of 3D stimuli can differ enough to yield significantly different discrimination performance [35]. In order to understand how the visual system integrates these three sources of information, we modeled different neural architectures and compared the model results to the observed data. If the visual system represented all three cues independently, then the precision of 3D perception could be greater than observed in this study, and elsewhere [3–5]. This raises two questions. First, what neural architecture can account for the observed perceptual results? Second, given the individual cue sensitivities, how close does the visual system come to maximizing the precision of 3D perception for combined-cue stimuli?

To address these questions, we used Bayesian decoding of simulated neuronal population responses [10]. Responses to stereoscopic, left eye perspective, and right eye perspective cues were simulated for each monkey based on recordings from neurons in the caudal intraparietal (CIP) area of the same animal (see **Methods**). Area CIP neurons are implicated in 3D perception since they are selective for 3D surface orientation, and their activity functionally correlates with behavioral reports of 3D orientation [32, 36–40]. We tested three neural architectures for combining cue-isolated responses (Fig 7A), and decoded the resulting combined-cue representation to simulate perceptual data (Fig 7B). Since the precision (but not the accuracy) of the simulated perceptual data depended on the architecture, we compared the decoded model precisions to the observed precisions in the monkey data (Fig 7C).

**Fig 7.**
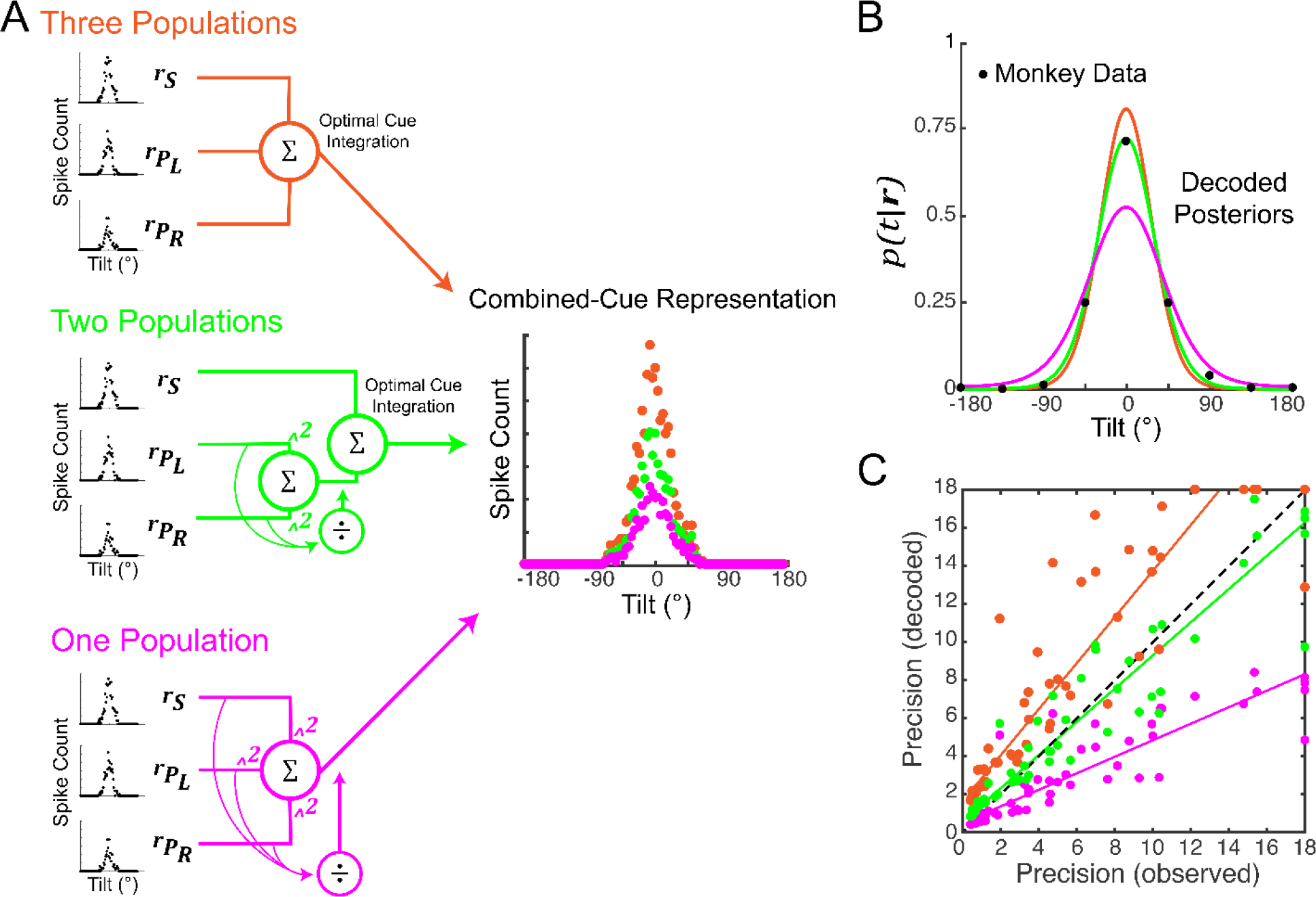
Cue integration is optimized but not maximized. **(A)** Schematics of three possible neural architectures for integrating population responses to stereoscopic cues (***r***_*s*_), left eye perspective cues 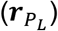, and right eye perspective cues 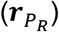. Top: Three independent populations represent each cue (orange). Middle: Two independent populations represent stereoscopic cues and perspective cues regardless of the stimulated eye (green). Bottom: One population estimates tilt using all three cues (magenta). Resulting combined-cue representations for each architecture are shown to the right. **(B)** Tilt posteriors, *p*(*t*|***r***), decoded from the combined-cue representations. Black dots show corresponding data from Monkey L. Given the same cue-isolated responses, precision was greatest for the three population model and lowest for the one population model. Note the close correspondence between the monkey data and the two population model’s decoded posterior. **(C)** Comparison of decoded model precisions and observed monkey precisions. Each point shows precision for a single pose. The three population model was more precise than the monkeys (nearly all points are above the identity line, black dashed). The two population model closely matched the monkeys’ performance (points are distributed about the identity line). The one population model was less precise than the monkeys (nearly all points are below the identity line). Solid lines show type-II regressions (*κ* = 18 excluded).

The first architecture assumed three independent neuronal populations, each of which represents tilt based on one of the three cues. Optimal integration is achieved by summing the three population responses [10] (Fig 7A, top). When we compared the monkey and decoded precisions, we found that the model was significantly more sensitive than the monkeys, Wilcoxon signed-rank test, *p* = 4.5×10^−9^ (Fig 7C, orange points). Indeed, if the three cues were represented independently and optimally integrated, the precision of combined-cue perception would have been, on average, 2.04 times greater than observed (ratio of decoded/observed precisions). This confirms our hypothesis that 3D tilt perception is less precise than theoretically possible with three independent cue representations. However, it is possible that observed tilt perception results from optimal integration of non-independent neuronal representations [3].

The second architecture tested this possibility, and assumed two independent populations that represent tilt based on either stereoscopic cues or perspective cues (regardless of the stimulated eye). For the perspective cue population, when both eyes are stimulated, the left and right eye driven responses are combined with a quadratic nonlinearity and divisively normalized (Fig 7A, middle). A similar model describes V1 responses to compound stimuli, and the operations combining the responses are widely implicated in neural processing [11–17]. As a consequence of divisive normalization, the independence of the two perspective cue representations is reduced, thereby decreasing the improvement in perceptual precision that results from having two cues. Optimal integration with this architecture is achieved by summing the stereoscopic and perspective cue population responses. When we compared the observed and decoded precisions (Fig 7C, green points), we found that they were not significantly different (Wilcoxon signed-rank test, *p* = 0.25). Thus, the perceptual results are consistent with a neural architecture in which two independent populations represent stereoscopic cues and perspective cues (from both eyes, combined using nonlinear canonical neural computations).

Lastly, we considered the possibility that a single neuronal population estimates tilt from both stereoscopic and perspective cues. When both eyes are stimulated, responses driven by each of the three cues are combined with a quadratic nonlinearity and divisively normalized (Fig 7A, bottom). As such, none of the cues are represented independently, and no explicit cue integration is required. When we compared the observed and decoded precisions, we found that the model was significantly less sensitive than the monkeys, Wilcoxon signed-rank test, *p* = 4.5×10^−9^(Fig 7C, magenta points).

These results identify a plausible neural architecture that can account for perceptual cue integration findings in both humans and monkeys, and rule out alternatives. They further identify the processing of left and right eye perspective cues within a single neuronal population as a potential factor limiting the precision of 3D perception, and demonstrate that 3D perception is optimized but not maximized as a result of canonical neural computations.

## Discussion

We evaluated the contributions of stereoscopic and perspective cues to 3D perception in macaque monkeys. Since the reliability of 3D cues is strongly affected by changes in depth or slant (Fig 1), we used an eight-alternative forced choice tilt discrimination task as a proxy for estimating how 3D sensitivity depends on object pose (i.e., orientation and position). We found that 3D perception was generally accurate across a wide range of poses. Instances of poor accuracy were largely restricted to stereoscopic cue stimuli with particularly low cue reliability. Thus, poor accuracy presumably reflected low certainty about the plane’s tilt and difficulty performing the task. Precision showed a clear pose dependence. For stereoscopic cues, precision increased with slant and decreased with distance from the fixation plane. For perspective cues, precision increased with slant and was independent of distance. At large distances and small slants, perspective cues were the sole contributor to 3D tilt perception, indicating that perspective cues extend 3D perception beyond the range supported by stereopsis.

### Evidence for a 3D oblique effect

The oblique effect for 2D tilt is characterized by larger biases and lower precisions at oblique compared to cardinal tilts [23–25, 27], and thought to reflect a prior for natural scene statistics [41]. A recent human study similarly found an oblique effect for 3D tilt with natural scene patches [27], consistent with a prior for the statistics of planar tilt [28, 29]. We examined if monkeys show an oblique effect for planar tilt, while testing for individual differences and cue-specific dependencies. Monkey L had larger biases at oblique than cardinal tilts in all three cue conditions, and lower precision at oblique than cardinal tilts in the combined-cue and perspective cue conditions. Such tilt dependencies were not as evident in Monkey F, indicating individual differences in the 3D oblique effect, similar to those for the 2D oblique effect in humans [24, 25]. Given the extensive training with both cardinal and oblique tilts, it is unlikely that training accounts for the oblique effect in Monkey L. Furthermore, both monkeys showed systematic biases with stereoscopic cue stimuli when precision was low, such that their reports were pulled towards ‘bottom-near’. This is consistent with the influence of a prior for ground planes, which occur in preponderance in natural scenes [28, 29]. Thus, Monkey F may have a weaker prior than Monkey L, such that only the ground plane component had an observable impact on perception. It is unlikely that the bottom-near bias reflected a preference for making downward saccades since horizontal eye movements are more accurate than vertical eye movements in humans [42, 43], and the oculomotor systems of macaques and humans are highly similar [44]. The results thus suggest that 3D perception in both humans and non-human primates is influenced by a prior for the 3D orientation statistics of natural scenes, and that the strength of that influence differs across individuals.

### Optimal cue integration

We used cue integration theory to predict combined-cue performance from stereoscopic and perspective (left and right eyes pooled) cue performances, and found that the cues were optimally integrated to achieve robust 3D perception. While this is consistent with previous human studies [4, 5], it is somewhat surprising since the theory assumes independent cue representations, but complete independence is unlikely (e.g., due to common retinal processing) [3]. The finding thus implies that the major sources of noise in 3D tilt estimation based on stereoscopic and perspective cues are largely independent. This could occur if the two estimates are created within different neuronal populations. Since the stimuli were interleaved, our finding of an optimal integration strategy further implies that the cues are dynamically reweighted to match the vagaries of cue reliabilities that occur with moment-to-moment changes in viewing conditions, such as happens every time the eyes move. Together with previous human studies [3–9], the current findings suggest that reliability-dependent cue integration is a conserved mechanism by which primates achieve robust 3D vision, and validate the macaque monkey as an ideal model system for studying the neural basis of 3D cue integration.

### Canonical computations optimize, but do not maximize 3D perception

We found that the observed perceptual results were optimal for a neural architecture in which two independent populations represent 3D tilt based on stereoscopic cues and perspective cues. To account for the perceptual results, it was essential that left and right eye perspective cue responses be combined with a quadratic nonlinearity and divisively normalized. Due to divisive normalization, the contributions of the two eyes’ perspective cues to perception will range from averaging (both signals contribute equally) when they are equally reliable to winner-take-all (only the more reliable signal contributes) when they differ substantially [11]. Thus, the model accounts for previous human cue integration findings showing cue averaging for balanced perspective cues [4, 5], and winner-take-all behaviors for imbalanced perspective cues [35]. Since divisive normalization reduces the independence of the two perspective cue responses, the computation imposes limits on the precision of perception. Our simulations showed that independent representations of stereoscopic, left eye perspective, and right eye perspective cues would double the precision of 3D perception.

Why would evolution not select for an integration strategy with higher precision? One possibility is the biological inefficiency associated with the sheer number of neurons required to maintain three independent cue representations, and the duplication of computational units to separately estimate 3D information from left and right eye signals. Another possibility is that 3D estimates derived from perspective cues are noisy, and combining left and right eye signals with divisive normalization attenuates that noise. The result suggests that 3D perception is optimized (through linear combinations of independent stereoscopic and perspective cue population responses), but is not maximized (due to divisive normalization of left and right eye perspective signals). We are currently testing this hypothesis with electrophysiological studies in the same monkeys. The results further serve as a reminder that “optimal” is in the eye of the beholder, and is most meaningful in the context of a specific neural architecture. We predict that analogous processes exist in other sensory systems which have multiple inputs sensitive to the same signals, as occurs in audition, vestibular processing, and bimanual touch.

## Methods

### Subjects and preparation

All surgeries and experimental procedures were approved by the Institutional Animal Care and Use Committee (IACUC) at the University of Wisconsin–Madison (Protocol G005229), and were in accordance with the National Institutes of Health’s Guide for the Care and Use of Laboratory Animals. All efforts were taken to ensure the well-being of the animals, including daily enrichment. Two male rhesus monkeys (*Macaca mulatta*) participated (Monkey L: 5 years of age, ~7.8 kg in weight; Monkey F: 4 years of age, ~5.5 kg in weight). A Delrin ring for stabilizing the head during training and experimental sessions was attached to the skull under general anesthesia [32, 38, 39]. After recovery, each monkey was trained to sit in a custom primate chair with head restraint, and to fixate a visual target within 2° version and 1° vergence windows for a liquid reward. We verified the ability to perceive stereoscopically-defined depth by having the monkeys fixate simulated targets between −20 and 40 cm of depth from the screen [45]. Binocular eye position was monitored optically at a sampling rate of 1,000 Hz (EyeLink 1000 plus, SR Research).

### Experimental control and stimulus presentation

Experimental control was performed using an open-source, network-based parallel processing framework [45]. Stimuli were created in MATLAB using Psychtoolbox 3 [46], and rendered using an NVIDIA GeForce GTX 970 graphics card on a Linux workstation (Ubuntu 16.04 LTS, Intel Xeon Processor, 24 GB RAM). A DLP LED projector (VPixx Technologies, Inc.) was used to rear project the stimuli onto a polarization preserving screen (Stewart Film Screen, Inc.). Stimuli were projected at 1,280 × 720 pixel resolution with a 240 Hz refresh rate. The screen distance was ~57 cm. The projected area subtended ~70° × 43° of visual angle. Stereoscopic presentation was achieved by sequencing the presentation of stimulus ‘half-images’ to each eye (120 Hz/eye) using a circular polarizer synchronized to the projector. Polarized glasses were worn.

### Visual stimuli

Planar surfaces were defined using random dot patterns (N = 250 dots). At the plane of fixation, dots subtended 0.35° of visual angle. The dots were bright (37.8 cd/m^2^) on a gray (12.3 cd/m^2^) background, measured through the polarized glasses (PR-524 LiteMate, Photo Research). On the screen, the stimuli were circular in shape and subtended 20° of visual angle.

Planes were presented at all combinations of eight tilts (0° to 315° in 45° steps), four slants (15° to 60° in 15° steps), and either six (Monkey L: 37, 57, 77, 87, 107, and 137 cm) or eight (Monkey F: 37, 57, 77, 87, 97, 107, 117, and 137 cm) distances. At 37 cm, all dots were in front of the plane of fixation. At 57 cm, dots were distributed in front of and behind the plane of fixation. At 77 cm and beyond, all dots were behind the plane of fixation. Presenting the stimuli at distances where the dots were entirely in front of, distributed about, or entirely behind the plane of fixation prevented the monkeys from relying on local absolute disparity cues to perform the task, ensuring that they judged the tilt of the plane [5, 40]. Under natural viewing conditions, changes in slant or distance affect the retinotopic area subtended by an object. To not confound 2D retinal features and 3D structure, we held the retinotopic area constant for all stimuli [32, 38, 39].

Stimuli were defined by both stereoscopic and perspective cues (‘combined-cue’; Fig 2B and 2C), stereoscopic cues (Fig 2D), or perspective cues (Fig 2E). Combined-cue stimuli had a uniform distribution of dots across the plane. Left and right eye half images were rendered by using perspective geometry to project each dot onto the appropriate screen position for the given eye. The perspective cues thus included retinal density gradients, foreshortening, and scaling. To ensure that the perspective cues only provided orientation information [5], the dots were scaled according to the plane’s distance such that their screen size depended only on the slant and tilt. Stereoscopic cue stimuli were created by defining a uniform distribution of dots on the screen and using ray tracing to assign each dot to a location on the plane. All dots had a circular shape and subtended 0.35°, irrespective of the pose. As such, the stereoscopic cue stimuli were designed to not contain any perspective cues that could be used to judge orientation. This was verified perceptually (S4A Fig). The combined-cue and stereoscopic cue stimuli were presented to both eyes. The perspective cue stimuli were the same as the combined-cue stimuli, but only one eye saw the planar stimulus (pseudo-randomly selected each trial) to eliminate stereoscopic cues. Both eyes saw the fixation target.

### Tilt discrimination task

The monkeys were trained to discriminate planar tilt in an 8AFC task. Task training began once a monkey could fixate a target on a blank screen for 2 s. They first learned to perform a two-alternative (right-near vs. left-near) task with all cue conditions and distances interleaved. The correct choice target was initially presented at a higher contrast than the distractor, and the contrast difference was reduced with training. Once an 80% correct rate with equal target contrasts was reached, all four cardinal tilts were introduced with a target contrast difference. Once a 50% correct rate was reached with equal contrasts, we started alternating training days between four cardinal and four oblique tilts. Once a 70% accuracy rate was reached with both cardinal and oblique tilts, all eight tilts were introduced together. Data collection began after performance in the 8AFC task stabilized.

In the task (Fig 2A), a monkey first acquired fixation of a target at the center of an otherwise blank screen. The target was a red circular dot (10.6 cd/m^2^ through the polarized glasses) subtending 0.3° of visual angle. After fixating for 300 ms, a plane centered on the target was presented for 1,000 ms. Fixation was then held for an additional 500 to 1,500 ms (pseudo-random duration) with no plane present. The fixation target then disappeared, and eight choice targets appeared at an eccentricity of 11° with polar angles ranging from 0° to 315° in 45° steps. The side of the plane nearest the monkey was reported with a saccade to the choice target at the corresponding polar angle for a liquid reward. A trial was aborted if fixation was broken before the choice targets appeared or if a choice was not made within 500 ms of their appearance. During the task, version and vergence were enforced with 2° windows. Offline, we calculated the time-averaged vergence error during the stimulus presentation for each trial. A 1° vergence window was then used to eliminate trials with errors ≥ 0.5° in magnitude.

Stimuli were presented in a pseudo-random order using a block design. A block consisted of one repetition of each combination of tilt, slant, distance, and cue type (Monkey L: 576 trials/block; Monkey F: 768 trials/block). The data set included 23,942 trials for Monkey L and 55,726 trials for Monkey F.

### Analyses

To quantify performance, we computed probability density functions describing the errors in reported tilts as follows. First, we took the difference between the reported tilt and the presented tilt for each trial: ΔTilt = Reported Tilt − Presented Tilt. Second, we created an error distribution of reported tilts by calculating the probability that the monkey reported ΔTilt. This was performed separately for each combination of slant, distance, and cue type. Depending on the analysis, error distributions were calculated using data at: (*i*) one tilt, (*ii*) cardinal (0°, 90°, 180°, and 270°) or oblique (45°, 135°, 225°, and 315°) tilts, or (*iii*) all tilts. Third, a von Mises probability density function was fit to each error distribution using maximum likelihood estimation:

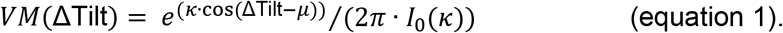

This function has two parameters: the mean (μ) and concentration (*κ*), which capture the accuracy and precision of perception, respectively. The closer μ is to 0, the more accurate (less biased) the judgments. The larger *κ*, the more concentrated (taller and narrower) the distribution, indicating more precise judgments. A modified Bessel function of order 0, *I*_0_(*κ*), normalizes the function to have unit area. The tilt sampling resolution limits the maximum *κ* that can be reliably estimated. We set an upper bound of *κ* = 18 in the maximum likelihood estimation routine, which corresponds to ~90% of the probability density function falling within the 45° tilt sampling interval.

To evaluate the integration of stereoscopic and perspective cues, we compared the observed combined-cue bias (*μ*_*c*_) and precision (*κ*_*c*_) to predictions derived from cue integration theory for circular variables [30]. The predictions were created using the stereoscopic and perspective cue biases (*μ*_*s*_ and *μ*_*p*_, respectively) and precisions (*κ*_*s*_ and *κ*_*p*_, respectively) taken from the von Mises fits. The optimal combined-cue parameters (bias: 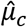; precision: 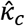) are:

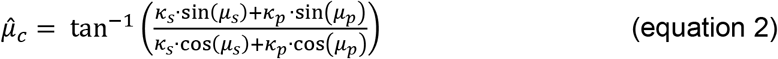

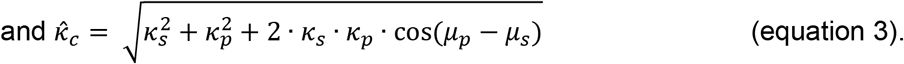

Circular statistics were used for μ [47]. Linear statistics were used for *κ*. All reported *p*-values are two-tailed. When multiple statistical comparisons were performed, *p*-values were adjusted using Bonferroni correction.

### Stereoscopic cue controls

We performed two controls to test if perception of the stereoscopic cue stimuli was affected by perspective cues. First, we tested if the stereoscopic cue stimuli contained perspective cues that could be used to perform the tilt discrimination task (S4A Fig). To elicit stereoscopic percepts, the stereoscopic cue stimuli were presented binocularly (both eyes saw the planar stimulus). To eliminate stereoscopic cues, they were presented monocularly (only one eye saw the planar surface, both eyes saw the fixation point). Above chance performance with monocularly viewed stimuli would indicate the presence of usable perspective cues. To maximize the potential perspective cue, the stimuli were presented at the largest tested slant (60°). They were presented at 57 cm. Parameters were otherwise the same as in the main experiment. All stimuli were presented interleaved. Monkey L completed 675 trials. Monkey F completed 1,819 trials.

Second, we considered if the stereoscopic cue precisions were affected by a potential stereoscopic–perspective cue conflict (S4B Fig). For the stereoscopic cue stimuli, the constant size, shape, and screen density of the dots can be interpreted as signaling zero slant. For stereoscopically defined non-zero slants, this could result in a perceived cue conflict which would increase with dot number since more isotropic dots provide more evidence of zero slant [5]. We therefore assessed precision with the stereoscopic cue stimuli as a function of dot number. A decrease in precision at larger dot numbers would indicate a cue conflict. To maximize the potential conflict, the stimuli were presented at the largest tested slant (60°). They were presented at 57 cm for Monkey L, and at 57 and 97 cm for Monkey F. Eleven dot numbers ranging from 5 to 250 (in steps of 25 starting at 25) were used. Parameters were otherwise the same as in the main experiment. All stimuli were presented interleaved. Monkey L completed 2,309 trials. Monkey F completed 5,844 trials at 57 cm, and 5,225 trials at 97 cm.

### Perspective cue control

We evaluated the perception of perspective cue stimuli after pooling responses to left and right eye stimulus presentations. To test the underlying assumption that perception was comparable for the two eyes, we independently fit von Mises probability density functions to the error distributions for each eye. Accuracies and precisions for the two eyes were then compared (S5 Fig).

### Neuronal models of cue integration

We used Bayesian decoding of model neuronal populations based on recordings from 3D surface orientation selective neurons in the caudal intraparietal (CIP) area to test if different neural architectures can account for the perceptual findings [10, 17]. Assuming independent neurons with Poisson spike count statistics, the probability that tilt (*t*) elicits population response (***r***) is:

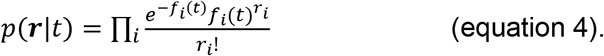

Here *f*_*i*_(*t*) and *r*_*i*_ are the *i*^th^ neuron’s tilt tuning curve and response, respectively. By Bayes rule and assuming a uniform prior, the posterior, *p*(*t*|***r***), describing the likelihood that *t* was presented given ***r*** is proportional to equation 4. As the number of neurons increases, *p*(*t*|***r***) converges to a Gaussian, and assuming *p*(*t*|***r***) guides behavior, the precision of perception (1/*σ*^2^ for that Gaussian) is proportional to the gain of the population activity (*g*). The constant of proportionality (*λ*) depends on the number of neurons and their tuning widths [10].

We simulated populations of monkey-specific neuronal tuning curves based on 3D surface orientation tuning curves measured in area CIP of the same animal (Monkeys L: N = 175; Monkey F: N = 94). The stimuli used in the neuronal recordings were the same as in the current study, except that only combined-cue stimuli were shown and the distances were 37, 57, 97, and 137 cm. We fit the tilt tuning curves at each slant–distance combination with a von Mises function [38]. Using these fits, we calculated the mean response amplitude and tuning width across neurons for each slant–distance combination and monkey, and linearly interpolated the values for untested distances. Using these parameters (DC offsets were not included in the model), we simulated 72 CIP neurons for each monkey, with 5° increments in tilt preference.

To determine the proportionality constant (*λ*) relating population gain to perceptual precision, we decoded the simulated population activity after scaling the responses by *λ*(*κ*_TW_), which depended on the pose-specific tuning width (*κ*_TW_) of the model neurons. We tried several functions to describe the relationship between *λ* and *κ*_TW_ (linear, exponential, double exponential, and two phase exponential). For each model, the parameters were fit to minimize the difference between the distributions of tilt errors made by the monkey and the decoded model posteriors. Fitting was performed separately for the two monkeys. Akaike’s Information Criterion was used to select the best fitting *λ* function. The best fit was provided by the exponential function (Monkey L: r = 0.89, *p* = 5.3×10^−7^; Monkey F: r = 0.82, *p* = 9.4×10^−7^), *λ* = *DC* + *G* · exp(*α* · *κ*_TW_), which was used in the simulations (Monkey L: *DC* = −1.6×10^−3^, *G* = 0.93, *α* = −1.87; Monkey F: *DC* = 2×10^−4^, *G* = 0.49, *α* = −1.89).

Next, we found response amplitudes for the cue-isolated conditions that minimized the difference between the distributions of tilt errors made by the monkey and the decoded model posteriors. For this, we assumed that the neuronal tuning widths did not differ across the visual cue conditions since CIP tilt tuning widths are similar regardless of the defining cue [36, 37]. The tilt tuning curves of the simulated neurons were thus:

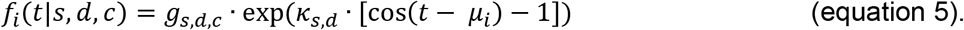

Here *f*_*i*_(*t*|*s*, *d*, *c*) is the *i*^th^ neuron’s tilt tuning curve for a given slant (*s*), distance (*d*), and visual cue (*c*). The gain (*g*_*s*,*d*,*c*_) depended on slant, distance, and visual cue. The tuning width (*κ*_*s*,*d*_) depended on slant and distance. The preferred tilt is *μ*_*i*_.

The model tuning curves were used to simulate neuronal population responses with Poisson variability for the cue-isolated conditions: stereoscopic (***r***_*s*_), left eye perspective 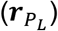, and right eye perspective 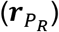. We tested three strategies for integrating these responses to create a combined-cue representation (***r***_*c*_) that was decoded after scaling the responses by *λ*(*κ*_TW_), see Fig 7. Since equation 4 converges to a Gaussian, the decoded posteriors were fit with Gaussian probability functions. To allow for a direct comparison, we refit the monkey tilt error distributions with Gaussians. Precision estimates from the von Mises and Gaussian fits were highly correlated (Monkey L: r = 1.0, *p* = 9.8×10^−25^; Monkey F: r = 0.99, *p* = 3.9×10^−26^).

#### Three populations

This model assumes three independent neuronal populations, each of which represents tilt based on a different cue: stereoscopic, left eye perspective, or right eye perspective. Optimal cue integration was achieved by summing the three cue-isolated population responses to create a combined-cue representation, ***r***_*c*_ [10]:

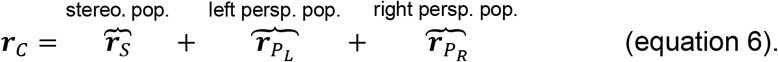

#### Two populations

This model assumes two independent neuronal populations which represent tilt based on stereoscopic cues or perspective cues from both eyes. The response of the perspective population to combined-cue stimuli is the divisively normalized sum of the squared single eye responses. Quadratic nonlinearities and divisive normalization are canonical computations that are broadly implicated in neural processing [11–17]. Optimal cue integration was achieved by summing the two cue-isolated population responses to create a combined-cue representation, ***r***_*c*_ [10]:

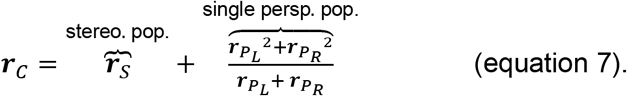

#### One population

This model assumes a single population of neurons that estimates tilt from both stereoscopic and perspective cues. The response of this population to combined-cue stimuli is the divisively normalized sum of the squared cue-isolated responses:

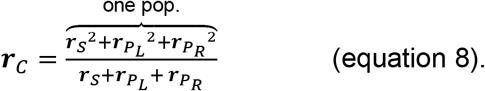

There are no free parameters in any of the models, so they can be directly compared. The responses of all three models to cue-isolated stimuli are equivalent, so distinguishing between them requires that their combined-cue predictions be compared to the performance of the monkeys.

## Acknowledgments

This work was supported by the Alfred P. Sloan Foundation FG-2016-6468, Whitehall Foundation Research Grant 2016-08-18, Shaw Scientist grant from the Greater Milwaukee Foundation, and National Institutes of Health Grant EY029438. Further support was provided by National Institutes of Health Grant P51OD011106 to the Wisconsin National Primate Research Center, University of Wisconsin – Madison.

## Author contributions

Conceptualization: TYC, BK, LT, AS, AR; Data Curation: TYC, BK, LT, RD, AR; Formal Analysis: TYC, LT, AR; Funding Acquisition: AR; Investigation: TYC, BK, LT, RD, AR; Methodology: TYC, BK, LT, AS, AR; Project Administration: AR; Resources: TYC, BK, LT, AS, RD, AR; Software: TYC, BK, LT, AS, RD, AR; Supervision: BK, AR; Validation: TYC, BK, RD, AR; Visualization: TYC, LT, AR; Writing – Original Draft Preparation: TYC, AS, AR; Writing – Review & Editing: TYC, LT, AS, AR.

## Supporting information

**S1 Fig.**
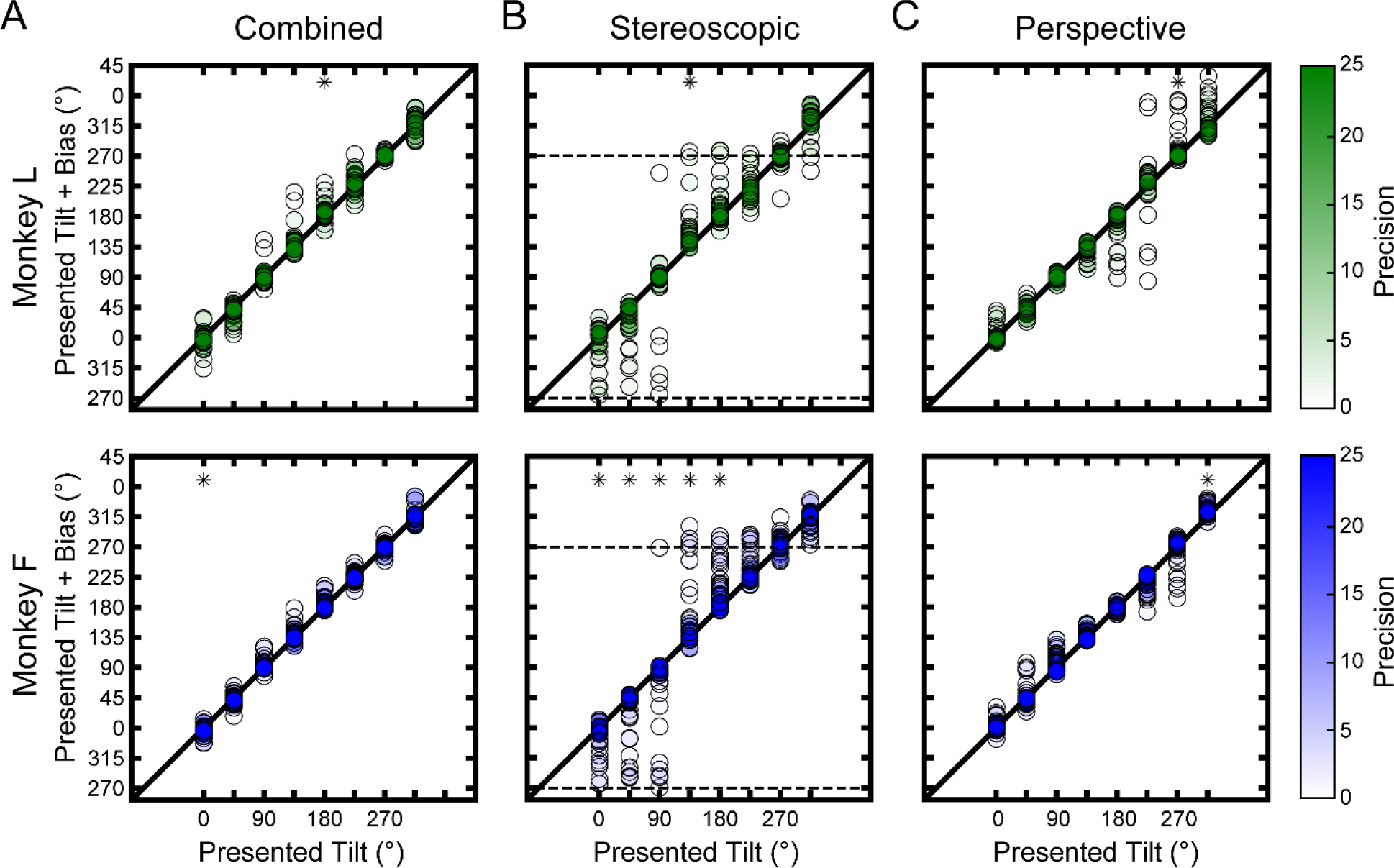
Bias as a function of tilt for each monkey and cue condition. The abscissa indicates the presented tilt and the ordinate indicates the presented tilt plus bias (*μ*). Black diagonals are identity lines. Greater vertical distance from identity indicates greater bias. Each point indicates the bias for a single pose. The fill opacity indicates the precision (*κ*). A circular median test was used to assess if the biases at each tilt were significantly different from 0°, corrected for multiple comparisons (N = 8 tilts). Asterisks mark significant biases. **(A)** Combined-cue stimuli. Significant biases occurred at 180° (median μ = 4.89°, *p* = 2.77×10^−4^) for Monkey L, and at 0° (median μ = −4.20°, *p* = 5.35×10^−4^) for Monkey F. In both cases, the median biases were small compared to the 45° tilt sampling interval. Absolute bias and precision were negatively correlated: Spearman r = −0.64, *p* = 3.20×10^−52^ (N = 448 slant × distance × tilt combinations, both monkeys). **(B)** Stereoscopic cue stimuli. Significant biases occurred at 135° (median μ = 16.03°, *p* = 2.77×10^−4^) for Monkey L, and at 0° (median μ = −6.40°, *p* = 2.10×10-3), 45° (median μ = −4.80°, *p* = 2.10×10^−3^), 90° (median μ = −7.61°, *p* = 2.10×10^−3^), 135° (median μ = 15.69°, *p* = 1.13×10^−4^), and 180° (median μ = 19.26°, *p* = 1.93×10^−5^) for Monkey F. Biases were most prevalent at low precisions, and the direction of the biases was consistent with perception being pulled towards a tilt of 270° (bottom-near). Absolute bias and precision were negatively correlated: Spearman r = −0.67, *p* = 1.29×10^−59^ (N = 448). **(C)** Perspective cue stimuli. Significant biases occurred at 270° (median μ = 3.07°, *p* = 1.54×10^−3^) for Monkey L, and at 315° (median μ = 8.92°, *p* = 2.46×10^−7^) for Monkey F. Absolute bias and precision were negatively correlated: Spearman r = −0.65, *p* = 1.79×10^−54^; N = 448).

**S2 Fig.**
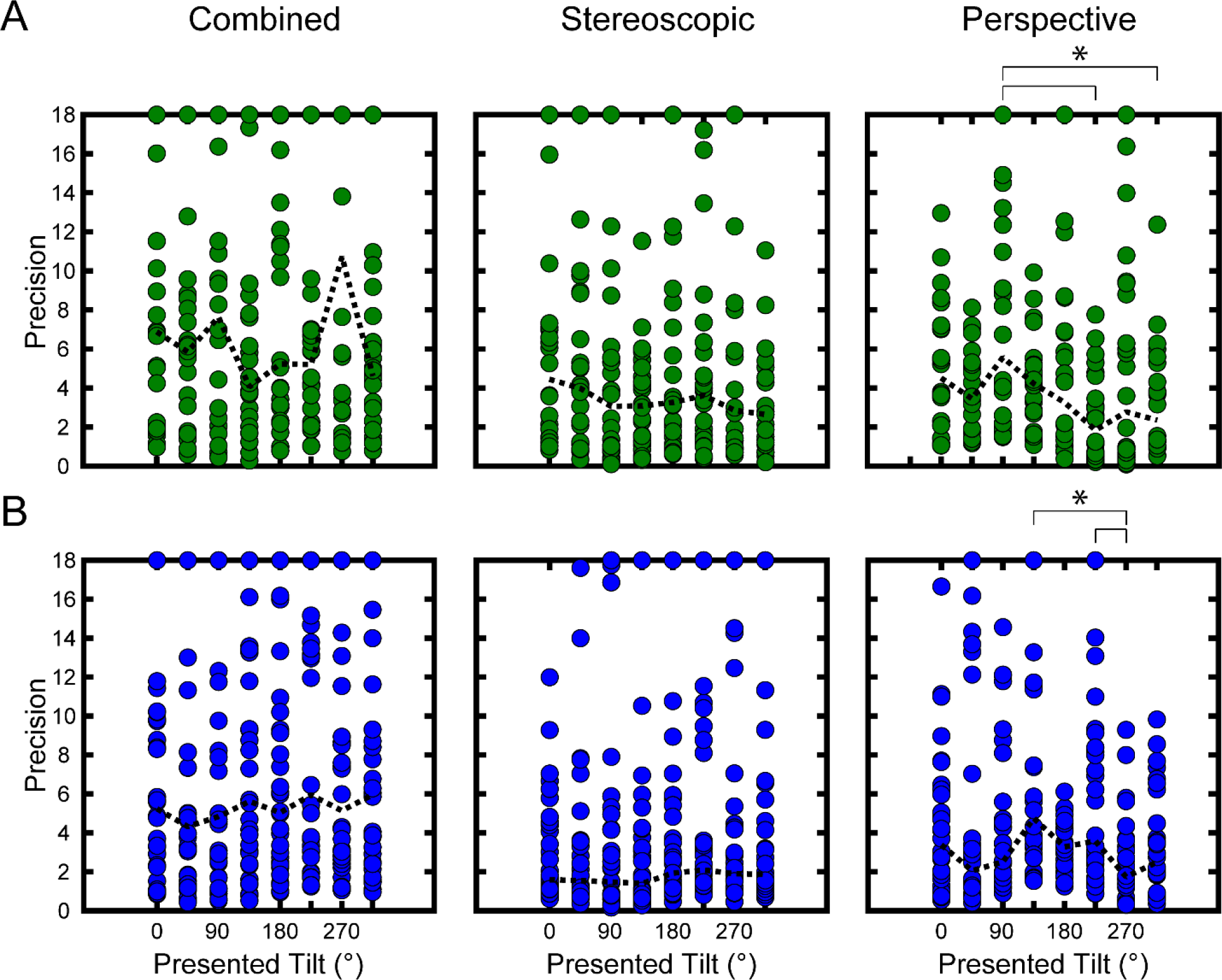
Precision as a function of tilt for each monkey and cue condition. The abscissa indicates the presented tilt and the ordinate indicates the precision of tilt perception (*κ*). Each point shows the precision for a single pose. Dashed black lines trace the median precisions. A Kruskal-Wallis test followed by Tukey’s honestly significant difference test was used to assess pairwise differences. Bracketed asterisks mark significant differences. **(A)** Combined-cue stimuli. Precision did not depend significantly on tilt (Monkey L: *p* = 0.15; Monkey F: *p* = 0.90). **(B)** Stereoscopic cue stimuli. Precision did not depend significantly on tilt (Monkey L: *p* = 0.69; Monkey F: *p* = 0.25). **(C)** Perspective cue stimuli. For Monkey L, the precision at 90° was significantly greater than at 225° (*p* = 9.27×10^−4^) and 315° (*p* = 0.04). For Monkey F, the precision at 270° was significantly lower than at 135° (*p* = 1.06×10^−3^) and 225° (*p* =0.02).

**S3 Fig.**
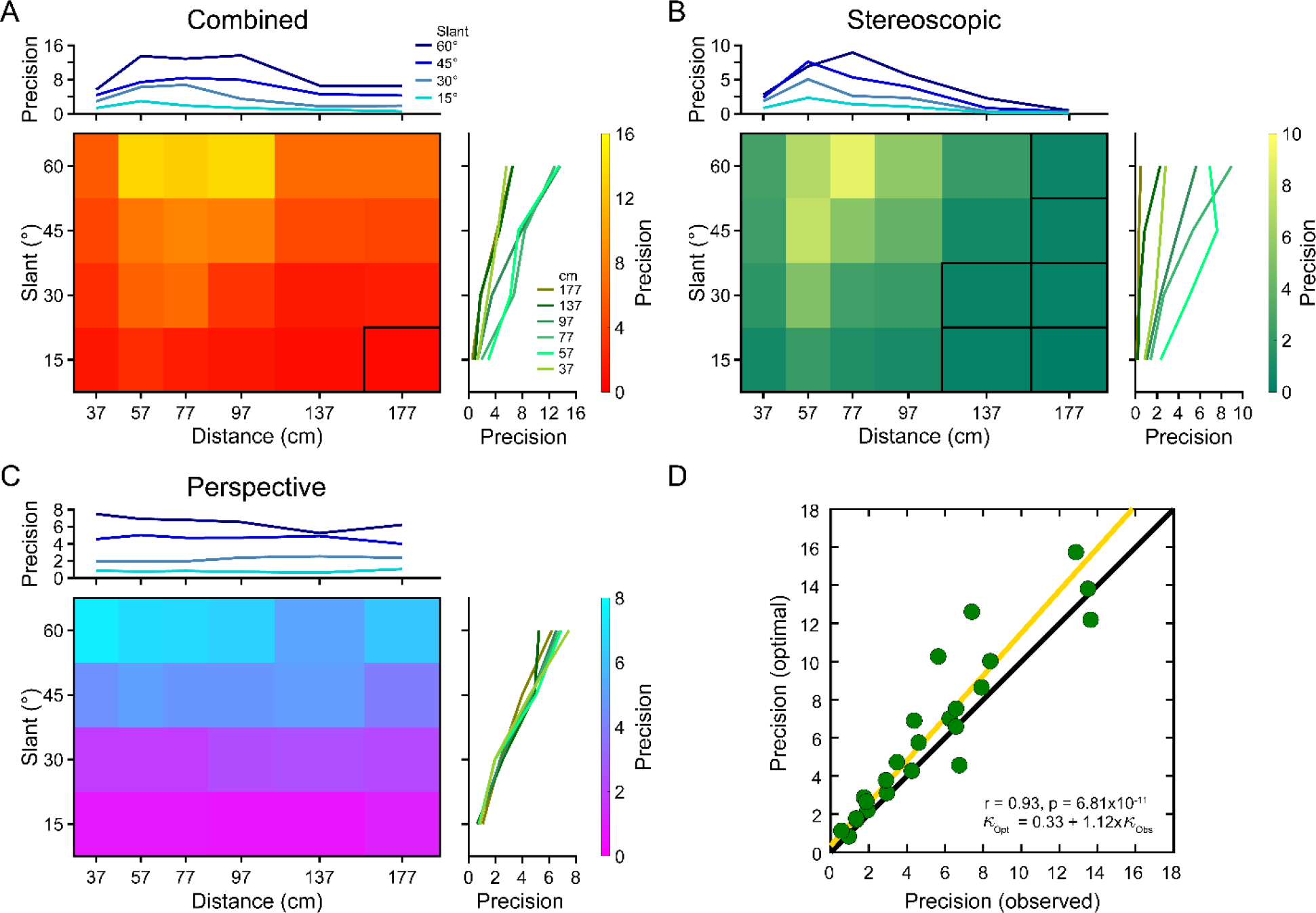
Precision and cue integration with larger stimuli. Monkey L also performed the tilt discrimination task with 30° stimuli defined by 550 dots. Four slants (15°, 30°, 45°, and 60°) and six distances (37, 57, 77, 97, 137, and 177 cm) were presented. A total of 7,508 trials were completed. **(A–C)** Heat maps showing precision (*κ*) as a function of slant and distance for each cue condition. Poses at which performance was not different from chance (Rayleigh test for circular uniformity, corrected for multiple comparisons) are outlined in black. **(A)** Combined-cue stimuli. **(B)** Stereoscopic cue stimuli. By 177 cm, performance was at chance levels for all slants. Perspective cue stimuli. **(D)** Cue integration. Each point shows the optimal vs. observed combined-cue precision (*κ*) for a single pose. The yellow line is the type-II regression line. Combined-cue precision was well predicted by optimal cue integration across a broad range of poses with different relative isolated-cue precisions.

**S4 Fig.**
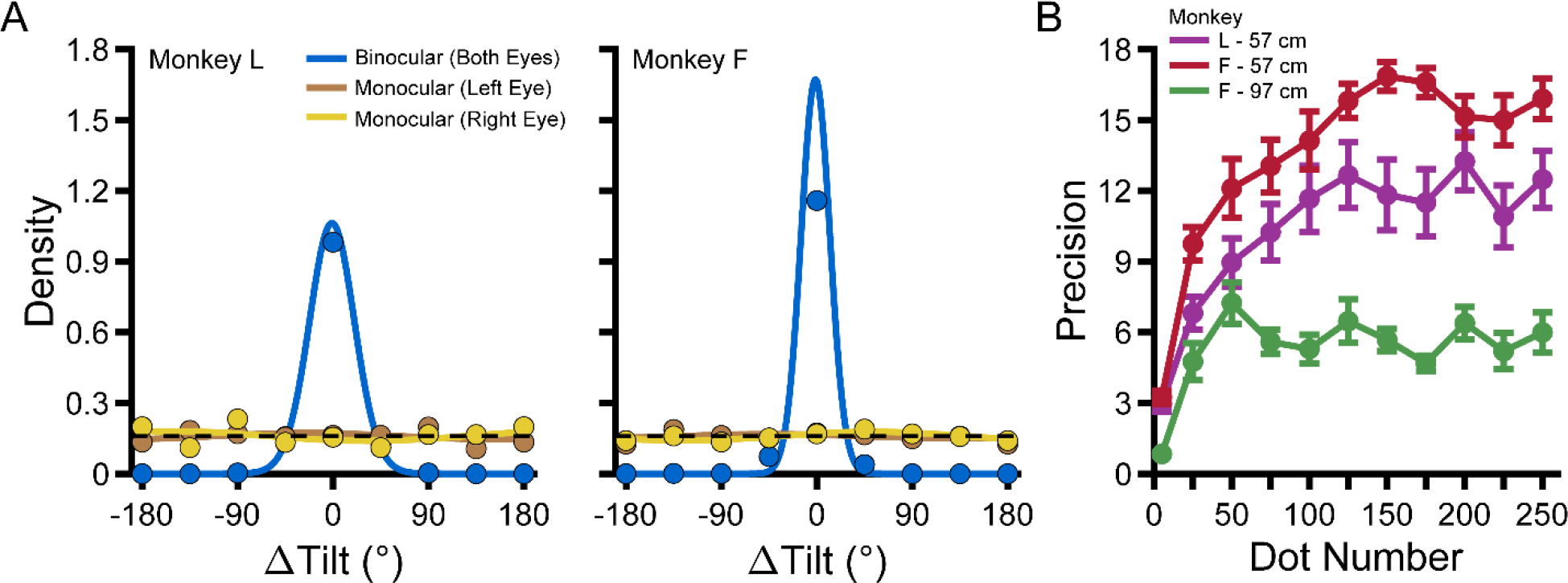
Stereoscopic cue controls. **(A)** Probability density functions with von Mises fits describing the errors in reported tilts made by Monkey L (left) and Monkey F (right) with stereoscopic cue stimuli, calculated using all 8 tilts. Stimuli were viewed binocularly (both eyes saw the planar stimulus; blue curves) or monocularly (one eye saw the planar stimulus; right eye stimulated: yellow; left eye stimulated: orange). Chance performance is indicated by the black dashed line. As expected, a Rayleigh test for circular uniformity confirmed significant binocular performance (Monkey L: *p* = 1.62−10^−230^; Monkey F: *p* = 2.23×10^−308^). In the monocular viewing conditions, performance was not significantly different from chance (Monkey L: left eye *p* = 0.72, right eye *p* = 0.62; Monkey F: left eye *p* = 0.69, right eye: *p* = 0.21). Thus, the stimuli contained no usable perspective information for performing the task. **(B)** Precision (*κ*) as a function of dot number, tested at 57 cm (Monkey L: purple; Monkey F: red) and 97 cm (Monkey F: green). Error bars show SEM across sessions. Precision depended significantly on dot number (Kruskal-Wallis test; Monkey L: *p* = 1.02×10^−7^; Monkey F: *p* = 3.82×10^−12^ at 57 cm; Monkey F: *p* = 8.71×10^−11^ at 97 cm). The initial increase with dot number was expected since more dots provide greater signal for performing the task. To test if any differences were the result of a decrease in precision, we ran pairwise comparisons using Tukey’s honestly significant difference test. In each case of a significant difference, the precision at the larger dot number was greater than at the smaller dot number. There were no significant differences between dot numbers ≥ 75. Thus, precision increased monotonically with dot number, suggesting that our stereoscopic cue precision estimates were not affected by a stereoscopic–perspective cue conflict.

**S5 Fig.**
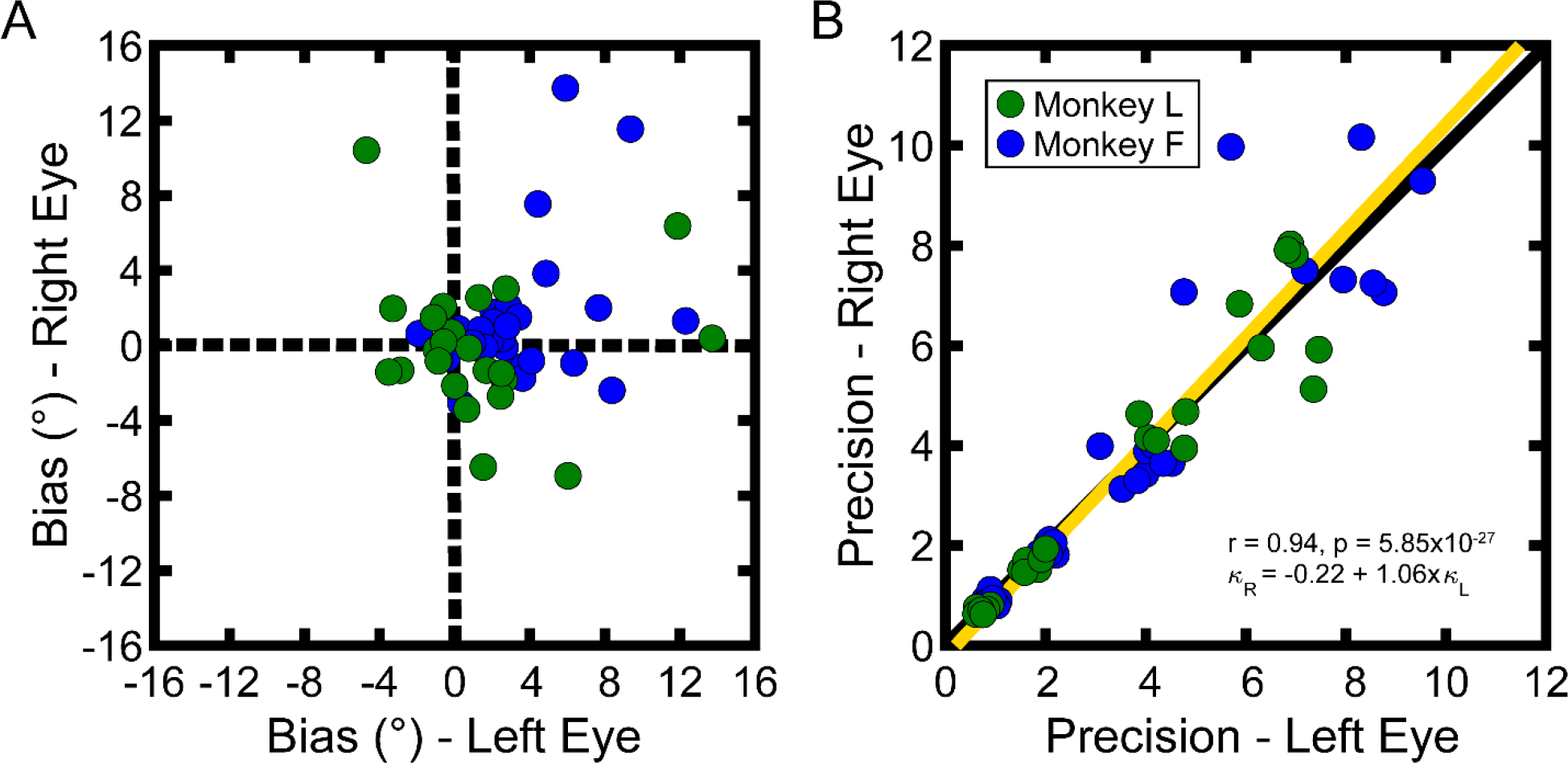
Perspective cue control. Each point corresponds to a single pose (Monkey L: N = 24; Monkey F: N = 32). **(A)** Right eye vs. left eye biases. Biases clustered near zero. The average difference between right and left eye biases was −2.12° for Monkey L and −1.87° for Monkey F. The difference was not significant for Monkey L (circular median test for multiple samples, *p* = 0.25), but was significant for Monkey F (*p* = 4.65×10^−4^). Although the difference was significant for Monkey F, it was less than for Monkey L and much smaller than the 45° tilt sampling interval. Thus, biases were small and comparable for the two eyes. **(B)** Right eye vs. left eye precisions. Across the two monkeys, the right and left eye precisions were highly correlated (r = 0.94, *p* = 5.85×10^−27^). The intercept (−0.22) and slope (1.06) of the type-II regression line (yellow) nearly specified the identity line (black diagonal). The precisions for the two eyes were not significantly different (Wilcoxon signed-rank test; Monkey L: *p* = 0.65; Monkey F: *p* = 0.16). The four clusters correspond to the four slants. Thus, precisions were comparable for the two eyes.

**S6 Fig.**
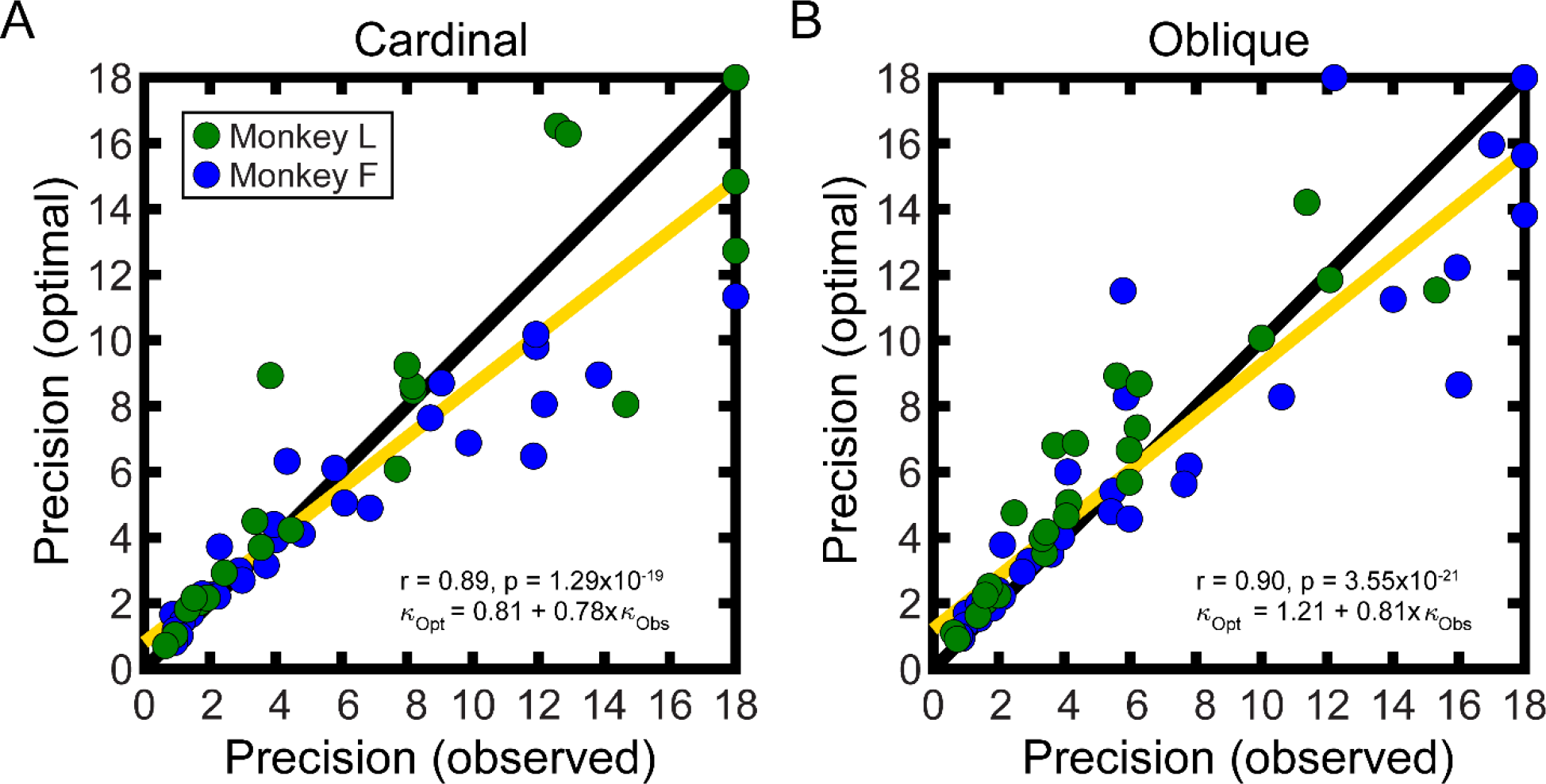
Cue integration at cardinal and oblique tilts. Each point shows the optimal vs. observed combined-cue precision for a single pose (Monkey L: N = 24; Monkey F: N = 32). Type-II regression lines are shown in yellow (*κ* = 18 excluded). **(A)** Cue integration at cardinal tilts. **(B)** Cue integration at oblique tilts.

## References

1. Barton RA. Binocularity and brain evolution in primates. Proc Natl Acad Sci U S A. 2004;101(27):10113–10115.

2. Hartley R, Zisserman A. Multiple view geometry in computer vision: Cambridge University Press; 2003.

3. Oruc I, Maloney LT, Landy MS. Weighted linear cue combination with possibly correlated error. Vision Res. 2003;43(23):2451–2468.

4. Knill DC, Saunders JA. Do humans optimally integrate stereo and texture information for judgments of surface slant? Vision Res. 2003;43(24):2539–2558.

5. Hillis JM, Watt SJ, Landy MS, Banks MS. Slant from texture and disparity cues: Optimal cue combination. J Vis. 2004;4(12):967–992.

6. Welchman AE, Deubelius A, Conrad V, Bulthoff HH, Kourtzi Z. 3D shape perception from combined depth cues in human visual cortex. Nat Neurosci. 2005;8(6):820–827.

7. Preston TJ, Kourtzi Z, Welchman AE. Adaptive estimation of three-dimensional structure in the human brain. J Neurosci. 2009;29(6):1688–1698.

8. Murphy AP, Ban H, Welchman AE. Integration of texture and disparity cues to surface slant in dorsal visual cortex. J Neurophysiol. 2013;110(1):190–203.

9. Rideaux R, Welchman AE. Proscription supports robust perceptual integration by suppression in human visual cortex. Nat Commun. 2018;9(1):1502.

10. Ma WJ, Beck JM, Latham PE, Pouget A. Bayesian inference with probabilistic population codes. Nat Neurosci. 2006;9(11):1432–1438.

11. Busse L, Wade AR, Carandini M. Representation of concurrent stimuli by population activity in visual cortex. Neuron. 2009;64(6):931–942.

12. Beck JM, Latham PE, Pouget A. Marginalization in neural circuits with divisive normalization. J Neurosci. 2011;31(43):15310–15319.

13. Carandini M, Heeger DJ. Normalization as a canonical neural computation. Nat Rev Neurosci. 2012;13(1):51–62.

14. Ni AM, Ray S, Maunsell JH. Tuned normalization explains the size of attention modulations. Neuron. 2012;73(4):803–813.

15. Louie K, Khaw MW, Glimcher PW. Normalization is a general neural mechanism for context-dependent decision making. Proc Natl Acad Sci U S A. 2013;110(15):6139–6144.

16. Qamar AT, Cotton RJ, George RG, Beck JM, Prezhdo E, Laudano A, et al. Trial-to-trial, uncertainty-based adjustment of decision boundaries in visual categorization. Proc Natl Acad Sci U S A. 2013;110(50):20332–20337.

17. Rosenberg A, Patterson JS, Angelaki DE. A computational perspective on autism. Proc Natl Acad Sci U S A. 2015;112(30):9158–9165.

18. Cormack R, Fox R. The computation of retinal disparity. Percept Psychophys. 1985;37(2):176–178.

19. Howard IP, Rogers BJ. Binocular vision and stereopsis: Oxford University Press, USA; 1995.

20. Stevens KA. The information content of texture gradients. Biol Cybern. 1981;42(2):95–105.

21. Knill DC. Surface orientation from texture: Ideal observers, generic observers and the information content of texture cues. Vision Res. 1998;38(11):1655–1682.

22. Prince SJ, Cumming BG, Parker AJ. Range and mechanism of encoding of horizontal disparity in macaque V1. J Neurophysiol. 2002;87(1):209–221.

23. Campbell FW, Kulikowski JJ, Levinson J. The effect of orientation on the visual resolution of gratings. J Physiol. 1966;187(2):427–436.

24. Westheimer G, Beard BL. Orientation dependency for foveal line stimuli: Detection and intensity discrimination, resolution, orientation discrimination and vernier acuity. Vision Res. 1998;38(8):1097–1103.

25. Girshick AR, Landy MS, Simoncelli EP. Cardinal rules: Visual orientation perception reflects knowledge of environmental statistics. Nat Neurosci. 2011;14(7):926–932.

26. Dakin CJ, Rosenberg A. Gravity estimation and verticality perception. Handb Clin Neurol. 2018;159:43–59.

27. Kim S, Burge J. The lawful imprecision of human surface tilt estimation in natural scenes. Elife. 2018;7.

28. Adams WJ, Elder JH, Graf EW, Leyland J, Lugtigheid AJ, Muryy A. The Southampton-York Natural Scenes (SYNS) dataset: Statistics of surface attitude. Sci Rep. 2016;6:35805.

29. Burge J, McCann BC, Geisler WS. Estimating 3D tilt from local image cues in natural scenes. J Vis. 2016;16(13):2.

30. Murray RF, Morgenstern Y. Cue combination on the circle and the sphere. J Vis. 2010;10(11):15.

31. Stevens KA. Slant-tilt: The visual encoding of surface orientation. Biol Cybern. 1983;46(3):183–195.

32. Rosenberg A, Cowan NJ, Angelaki DE. The visual representation of 3D object orientation in parietal cortex. J Neurosci. 2013;33(49):19352–19361.

33. Seilheimer RL, Rosenberg A, Angelaki DE. Models and processes of multisensory cue combination. Curr Opin Neurobiol. 2014;25:38–46.

34. Cormack LK, Czuba TB, Knoll J, Huk AC. Binocular Mechanisms of 3D Motion Processing. Annu Rev Vis Sci. 2017;3:297–318.

35. Thompson L, Ji M, Rokers B, Rosenberg A. Contributions of binocular and monocular cues to motion-in-depth perception. Journal of Vision. 2019;19(3):2.

36. Tsutsui K, Jiang M, Yara K, Sakata H, Taira M. Integration of perspective and disparity cues in surface-orientation-selective neurons of area CIP. J Neurophysiol. 2001;86(6):2856–2867.

37. Tsutsui K, Sakata H, Naganuma T, Taira M. Neural correlates for perception of 3D surface orientation from texture gradient. Science. 2002;298(5592):409–412.

38. Rosenberg A, Angelaki DE. Gravity influences the visual representation of object tilt in parietal cortex. J Neurosci. 2014;34(43):14170–14180.

39. Rosenberg A, Angelaki DE. Reliability-dependent contributions of visual orientation cues in parietal cortex. Proc Natl Acad Sci U S A. 2014;111(50):18043–18048.

40. Elmore LC, Rosenberg A, DeAngelis GC, Angelaki DE. Choice-related activity during visual slant discrimination in macaque CIP but not V3A. eNeuro. 2019;6(2):e0248.

41. Coppola DM, Purves HR, McCoy AN, Purves D. The distribution of oriented contours in the real world. Proc Natl Acad Sci U S A. 1998;95(7):4002–4006.

42. Kapoula Z, Yang Q, Vernet M, Dieudonne B, Greffard S, Verny M. Spread deficits in initiation, speed and accuracy of horizontal and vertical automatic saccades in dementia with lewy bodies. Front Neurol. 2010;1:138.

43. Ke SR, Lam J, Pai DK, Spering M. Directional asymmetries in human smooth pursuit eye movements. Invest Ophthalmol Vis Sci. 2013;54(6):4409–4421.

44. Fuchs AF. Saccadic and smooth pursuit eye movements in the monkey. J Physiol. 1967;191(3):609–631.

45. Kim B, Kenchappa SC, Sunkara A, Chang T-Y, Thompson L, Doudlah R, et al. Real-time experimental control using network-based parallel processing. eLIFE. 2019;8:e40231.

46. Kleiner M, Brainard D, Pelli D, Ingling A, Murray R, Broussard C. What’s new in Psychtoolbox-3. Perception. 2007;36(14):1.

47. Fisher NI. Statistical analysis of circular data: Cambridge University Press; 1995.

